# Increase in ER-mitochondria contacts and mitochondrial fusion are hallmarks of mitochondrial activation during embryogenesis

**DOI:** 10.1101/2024.06.11.598492

**Authors:** Anastasia Chugunova, Hannah Keresztes, Roksolana Kobylinska, Maria Novatchkova, Thomas Lendl, Marcus Strobl, Michael Schutzbier, Gerhard Dürnberger, Richard Imre, Elisabeth Roitinger, Pawel Pasierbek, Alberto Moreno Cencerrado, Marlene Brandstetter, Thomas Köcher, Benedikt Agerer, Jakob-Wendelin Genger, Andreas Bergthaler, Andrea Pauli

## Abstract

Mitochondrial ATP production is essential for development, yet the mechanisms underlying the continuous increase in mitochondrial activity during embryogenesis remain elusive. Using zebrafish as a model system for vertebrate development, we comprehensively profile mitochondrial activity, morphology, metabolome, proteome and phospho-proteome as well as respiratory chain enzymatic activity. Our data show that the increase in mitochondrial activity during embryogenesis does not require mitochondrial biogenesis, is not limited by metabolic substrates at early stages, and occurs without an increase in the abundance of respiratory chain complexes or their *in vitro* activity. Our analyses pinpoint a previously unexplored increase in mitochondrial-ER association during early stages in combination with changes in mitochondrial morphology at later stages as possible contributors to the rise in mitochondrial activity during embryogenesis. Overall, our systematic profiling of the molecular and morphological changes to mitochondria during embryogenesis provides a valuable resource for further studying mitochondrial function during embryogenesis.

## Introduction

Mitochondria play a crucial role in maintaining energy homeostasis, which is essential for fertilization and normal embryo development^1–6^. Over 90% of cellular ATP is generated within mitochondria through oxidative phosphorylation (OXPHOS), facilitated by five enzymatic complexes situated in cristae in the inner mitochondrial membrane. These include complexes I to IV, comprising the electron transport chain (ETC), and complex V (ATP synthase), which produces ATP by phosphorylation using the energy generated by the translocation of protons across the inner membrane by the ETC.

All mitochondria inherited by a new organism are maternally derived, which requires oocytes to store more mitochondria than any other cell type^7–9^. Mitochondria in mature oocytes display low mitochondrial activity^10–13^ and are characterized by a round shape and reduced number of cristae^14–16^. Inhibition of mitochondrial activity during oogenesis is thought to be achieved through the downregulation of certain respiratory chain complexes^12,13^ to mitigate oxidative stress that could damage oocytes and interfere with fertilization and/or subsequent embryo development^17^.

Mitochondria undergo an initial activation triggered by the release of calcium ions (Ca^2+^) from internal stores during fertilization and/or egg activation. Ca^2+^ stimulates Krebs cycle dehydrogenases^18–20^ and directly impacts the ATP synthase^21^, which results in a transient increase in cytosolic ATP levels. This early transient increase is essential for completion of meiosis^22–24^. As development progresses, mitochondrial activity was found to continuously increase across different species^11,25–28^. This increase was shown to occur in the absence of mitochondrial replication until blastocyst stages in mammals^29,30^, arguing against mitochondrial biogenesis as primary contributor to the observed increase in activity, at least during the early embryonic stages. Meanwhile, mitochondria were shown to mature during embryogenesis, regaining their conventional elongated shape and well-defined cristae^14–16^. However, apart from the correlation between changes in mitochondrial morphology and their activity, the mechanisms underlying the increase in mitochondrial activity during embryogenesis remain unclear, and the major changes in mitochondrial function remain poorly understood. In this regard, it is interesting to note that mitochondria are transcriptionally active already immediately after fertilization while nuclear transcription is still inactive^29,31,32^. This raises the possibility that mitochondrial activity may increase due to the synthesis of new respiratory chain complexes – an idea that has not yet been tested. In addition, it is unclear whether multiple, sequentially acting mechanisms may lead to the continuous increase in mitochondrial activity during embryogenesis.

In this study, we provide a comprehensive characterization of the molecular and morphological changes in mitochondria during embryogenesis, using zebrafish as a model system. We show that the initial rise in mitochondrial activity neither requires mitochondrial biogenesis nor changes in the abundance or enzymatic activity of respiratory chain complexes. Instead, we identify increased interaction between mitochondria and endoplasmic reticulum (ER) and elongation of mitochondria as hallmarks of mitochondrial activation during embryogenesis.

## Results

### Mitochondrial activity increases during embryogenesis independently of mitochondria biogenesis

To comprehensively characterize the molecular and morphological changes to mitochondria and the ETC (**Figure 1A**) during embryogenesis, we used zebrafish as a model system because the externally developing embryos provide easy access to samples for biochemical and functional studies at early embryonic stages. We first assessed the dynamics of mitochondrial activity in zebrafish embryos by measuring oxygen consumption of individual embryos over time. Respiration was low at early stages and progressively increased during embryogenesis (**Figure 1B**), which is consistent with published data from mammalian^11,25,27–29^ and zebrafish embryos^26,33,34^. Moreover, in agreement with prior studies in other organisms^29,30^ and in zebrafish^35,36^ showing the absence of mitochondrial biogenesis early on, we did not detect the mitochondrial DNA polymerase Polg by mass spectrometry in zebrafish embryos before one day post fertilization^37^. Consistently, we found that the mtDNA content per embryo remains constant for at least 14 hours post fertilization (hpf) showing that mitochondrial biogenesis does not contribute to the early increase in mitochondrial activity (**Figure 1C**).

**Figure 1.**
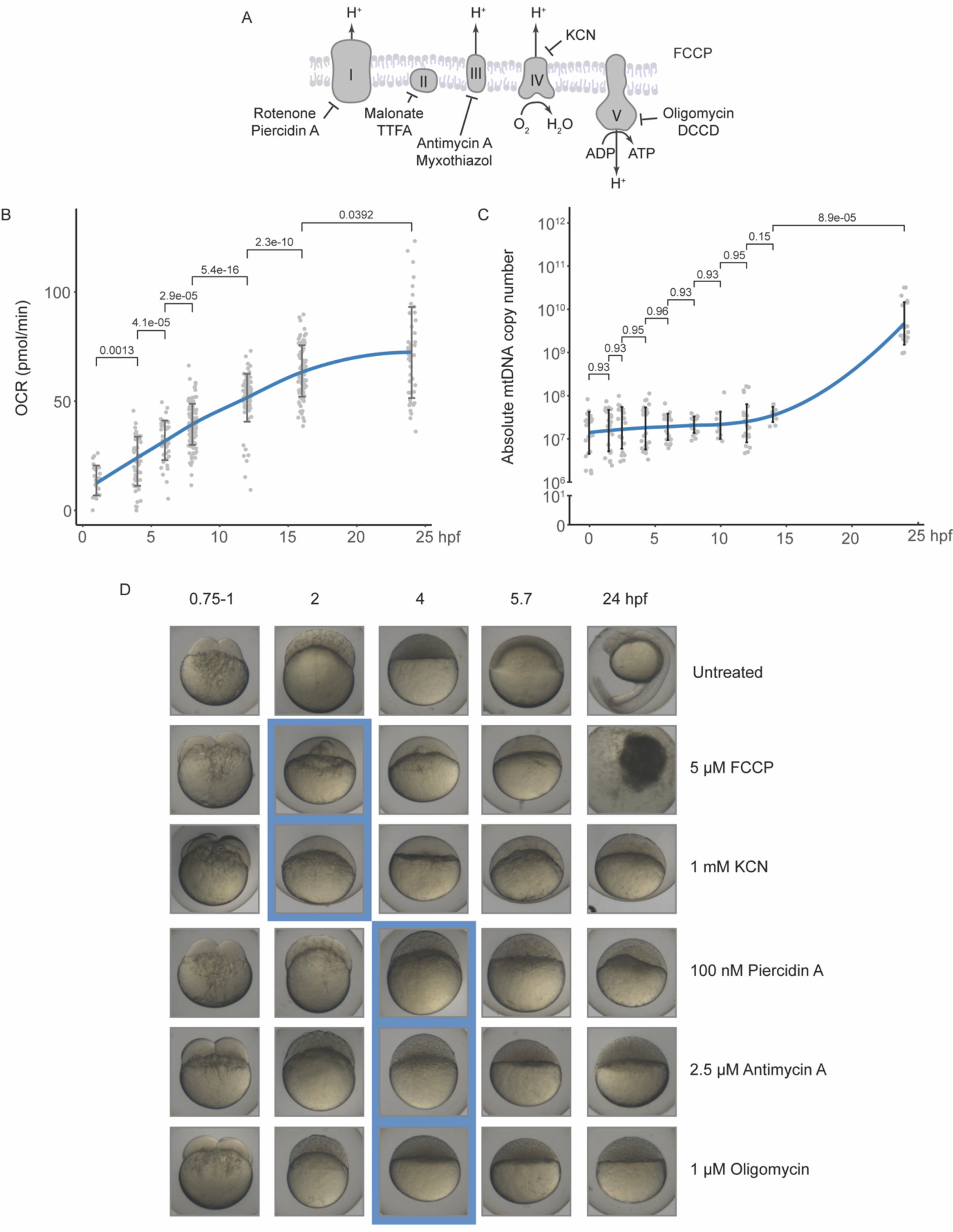
Mitochondrial activity is essential for early zebrafish embryogenesis, and it increases independently of mitochondrial biogenesis. (A) Scheme of the respiratory chain. Inhibitors blocking specific complexes are indicated. (B) Oxygen consumption rate (OCR; hpf stands for hours post fertilization) measured in individual zebrafish embryos at different developmental stages. Kruskal-Wallis One-Way ANOVA P-values (Benjamini-Hochberg correction) are indicated for statistically significant changes. The blue line represents the nonparametric regression with the standard deviation. (C) Absolute mtDNA copy number measured by RT-qPCR in individual zebrafish embryos. The blue line represents the nonparametric regression with the standard deviation. (D) Effects of respiratory chain inhibitors on zebrafish embryo development. The blue boxes indicate the stage when developmentally arrested embryos were observed.

To assess at which developmental stage mitochondrial activity is required for progression through embryogenesis in zebrafish, we treated embryos after fertilization with known inhibitors of each of the five complexes (**Figure 1A**). Since respiration is coupled to ATP production through the membrane potential, the increase in oxygen consumption during development could be necessary either for ATP production or for maintenance of the mitochondrial membrane potential, or for both. To test the importance of the ETC during early embryogenesis, we depolarized the inner mitochondrial membrane by treating embryos right after fertilization with FCCP (**Figure 1A, D** and **S1A, B**). To inhibit complex IV, we treated embryos with KCN (**Figure 1A, D** and **S1B**). Both treatments caused an almost immediate arrest in embryogenesis before the cleavage stages (**Figure 1D** and **S1B**). However, pharmacological inhibition of other electron transport chain complexes by treatment with Piercidin A (complex I), Antimycin A (complex III) or Oligomycin or DCCD (complex V) blocked progression through embryogenesis slightly later, yet still within the first hours of embryogenesis (**Figure 1A, D** and **S1B**). Together, these results indicate that the ETC activity is already required during the early stages of zebrafish embryo development.

Interestingly, embryos in which ATP production was inhibited by blocking complex V with Oligomycin treatment were unable to proceed beyond the sphere stage, the stage at which the major wave of zygotic genome activation (ZGA) takes place in zebrafish embryos^31,32^. However, RNA-sequencing revealed that ZGA occurs in Oligomycin-treated embryos, albeit at slightly reduced levels, in contrast to embryos in which transcription was inhibited by injection of the RNA Pol-II inhibitor Actinomycin-D (**Figure S1C**). This suggests that the reason for the developmental arrest upon inhibition of ATP production is not due to a complete failure to activate the zygotic genome.

Together, our data revealed that mitochondrial activity is essential for the early stages of zebrafish embryogenesis and progressively increases independently of mitochondrial biogenesis.

### Targeted metabolic profiling reveals remodeling of the metabolome in the early embryo

To identify possible contributors to the increase in mitochondrial activity during the early stages of embryogenesis, we performed a systematic analysis of metabolites and the mitochondrial proteome changes during the first day of embryogenesis. Given that the respiratory chain activity depends on NADH levels, we first tested whether substrate availability might be limiting in the early stages. Targeted detection of a specific set of metabolites and amino acids revealed dynamic metabolome remodeling (**Figure 2A** and **Figure S2A**). Most importantly, we found that the energy metabolites NADH, pyruvate and fructose-1,6-phosphate were abundant early on and subsequently utilized during development (**Figure 2A, B**). Conversely, the levels of dihydroxy-acetone-phosphate (DHAP), 3-phospho-glyceric acid (3PG) and phospho-enol-pyruvate (PEP), the metabolic intermediates of glycolysis, were initially low and increased after 6 hpf, indicating activation of glycolysis (**Figure 2A, B**). This was consistent with the increased abundance of glycolysis enzymes in the mitochondrial fraction during later embryonic stages (**Figure S2B**). Together with metabolic intermediates of glycolysis, a subset of Krebs cycle intermediates including succinate, malate, fumarate accumulated during gastrulation (**Figure 2A**) while the remaining Krebs cycle metabolites, namely citrate, aconitic acid, and α-ketoglutarate, increased, around 8 hpf. This imbalance in Krebs cycle metabolites could not be accounted for by differential expression of Krebs cycle enzymes, as their abundance remained unchanged during early embryogenesis (**Figure S2B**). Overall, our results in zebrafish embryos resemble the dynamics observed in mammalian preimplantation embryos, where glycolysis becomes active only at gastrulation^1,11^ and the pyruvate dehydrogenase complex in mitochondria^38^ is inactive, leading to the Krebs cycle imbalance.

**Figure 2.**
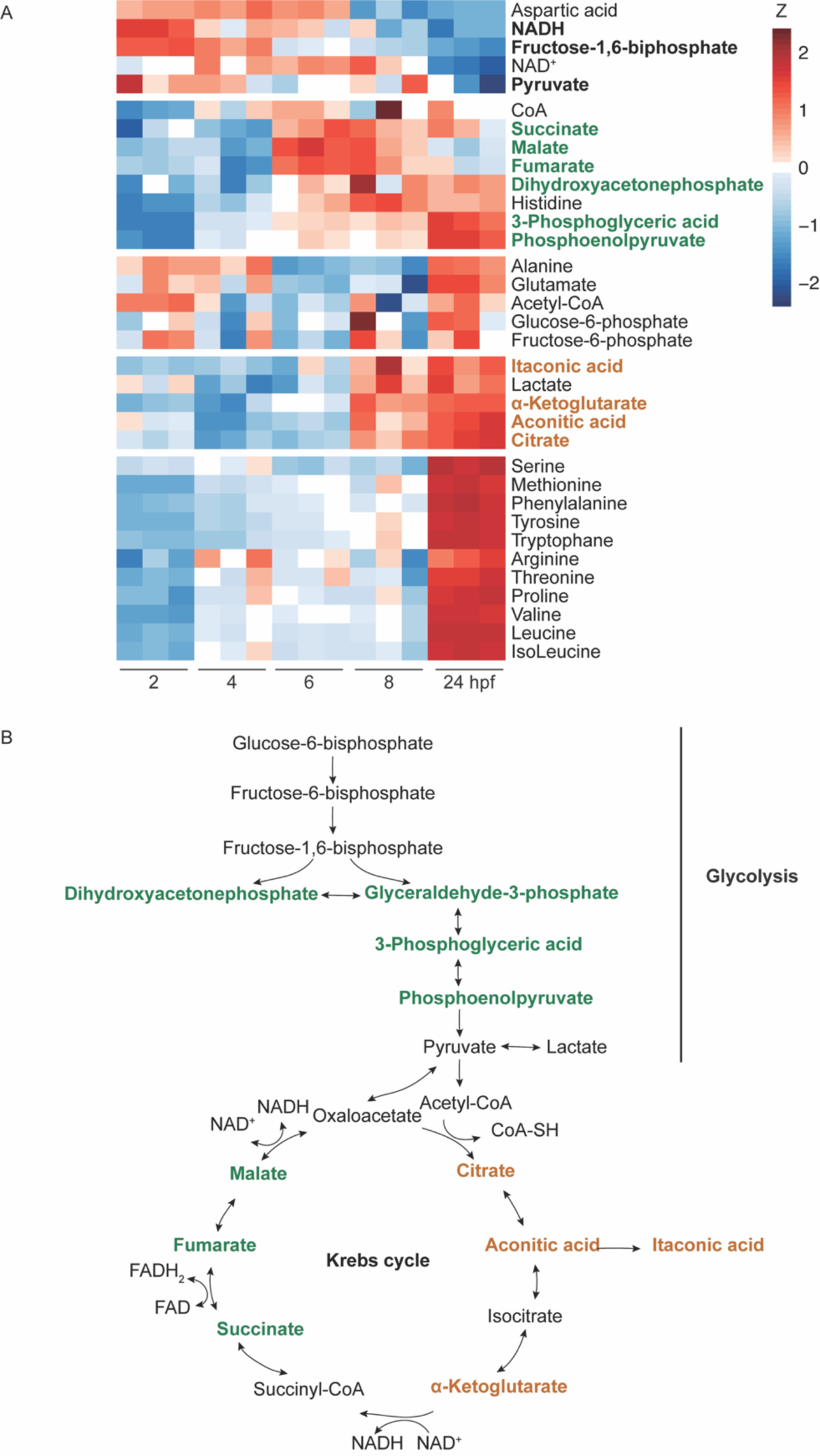
Respiratory chain substrates are present in the early zebrafish embryo and metabolites undergo major changes during embryogenesis. (A) Targeted metabolomics of zebrafish embryos at different developmental stages. Heatmap of a subset of metabolites that were measured during embryogenesis. The color code represents a z-score for the log-transformed raw values for each metabolite. Each row corresponds to an individual metabolite and one square is a single replicate (3 replicates for each stage). (B) Schematic representation of the glycolysis and Krebs cycle metabolites that are upregulated at 6 hpf (green) or at 8 hpf (orange).

Although NADH levels were high during early stages, it might not be available to mitochondria since intact mitochondria are impermeable to NADH^39^, and NADH transport across the mitochondrial membrane depends on the malate-aspartate shuttle^40^. However, malate-aspartate shuttle proteins were present and did not change significantly throughout the first 24 hours in our mitochondrial proteomics dataset (**Figure S2B**). Collectively, our results reveal that metabolites change dynamically during zebrafish embryogenesis. Moreover, NADH is present in early zebrafish embryos and gets utilized, ruling out substrate unavailability as reason for the low mitochondrial activity during the initial stages of embryogenesis.

### Abundance and *in vitro* activity of isolated respiratory chain complexes are constant during early embryogenesis

Mitochondrial activity could be regulated by increasing the abundance of respiratory chain complexes or by the formation of supercomplexes. For example, it has been reported that the downregulation of mitochondrial activity during oogenesis is achieved by the elimination of complex I in humans and frogs^12^, and by reduction in complex I and V in flies^13^. We therefore hypothesized that the increase in respiration could be due to an increase in the level and/or activity of respiratory chain complexes and supercomplexes. Assessment of the amount of respiratory chain complexes using blue native polyacrylamide gel electrophoresis (BNP) revealed no increase in the number of complexes and supercomplexes throughout the first 24 hours of zebrafish embryogenesis (**Figure 3A, B**), though we detected a slight yet reproducible reduction of complex I at 24 hpf (**Figure 3B**). Since previous studies have shown that different paralogues or modifications of respiratory chain subunits can modulate the activity of the electron transport chain in different tissues or biological processes^41–43^, we examined the expression levels of individual respiratory chain subunits over time using our mitochondrial proteomics data. In line with our analysis by BNP, expression levels of most of the ETC subunits were constant throughout embryogenesis (**Figure 3C**), and apart from an increase in protein expression of Ndufa4b, one of the paralogues of the complex IV subunit Ndufa4, and a decrease in complex V member Atp5mea at 24 hpf (**Figure 3C**), no other instances of differentially expressed subunits were observed. To rule out that these differentially expressed subunits or potential modifications to respiratory chain subunits could affect the activity of respiratory chain complexes during embryogenesis, we performed BNP followed by measurement of complex I, II, IV and V in-gel activity. The *in vitro* activity of complexes I, II, IV and V was found to be constant at different time points of embryogenesis (**Figure 3D-G**). We conclude that the expression levels of respiratory chain subunits remain constant during embryogenesis, and that the enzymatic activities of individual ETC complexes do not increase during early stages of embryogenesis, suggesting that other external factors might influence ETC activity *in vivo*.

**Figure 3.**
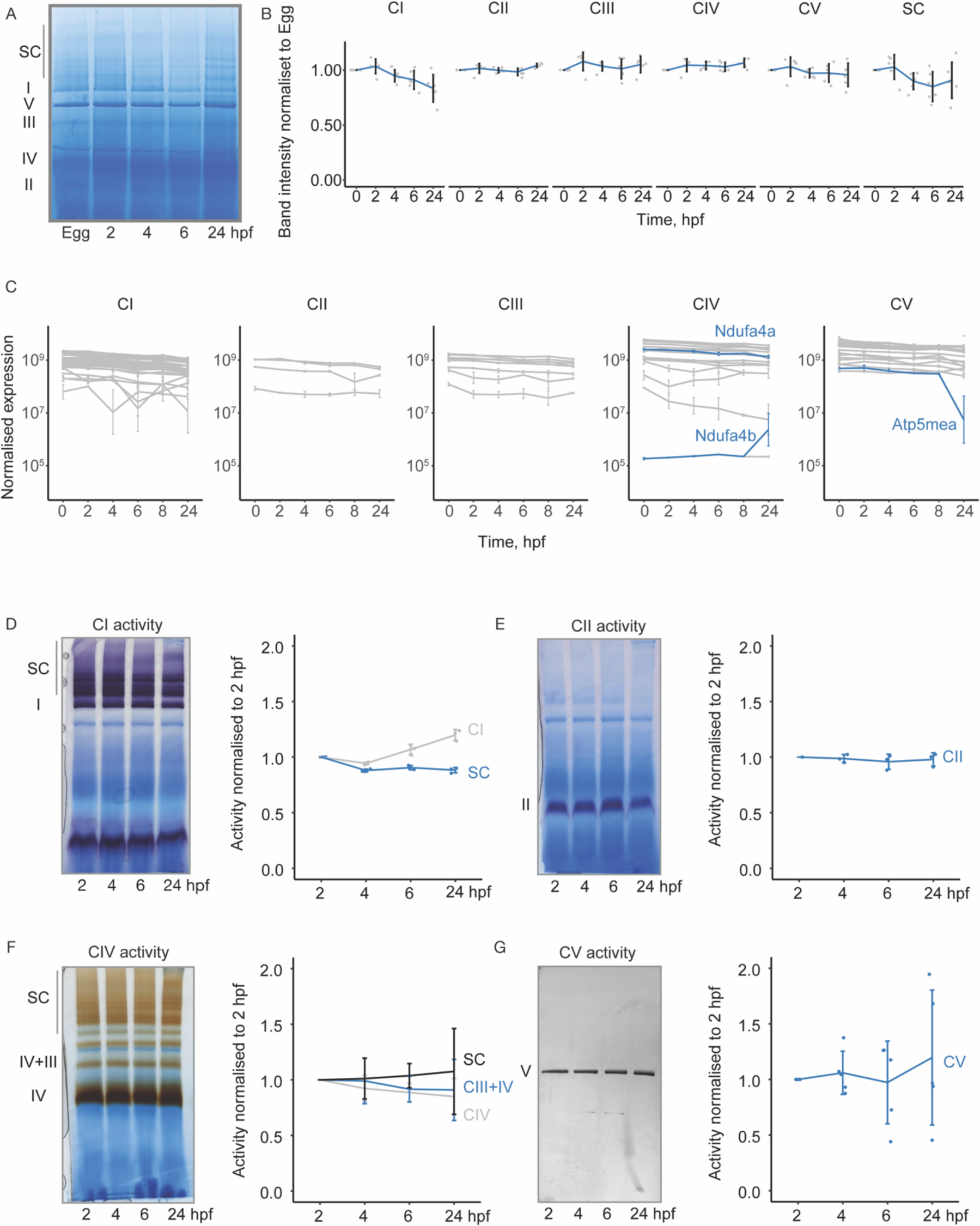
Abundances and enzymatic activities of respiratory chain complexes are constant during early zebrafish embryogenesis. (A) Blue native PAGE of digitonin-solubilized mitochondria isolated at different stages of zebrafish embryogenesis. Molecular weight is indicated on the left. SC stands for supercomplexes. (B) Quantification of the abundances of respiratory chain complexes at different developmental stages of zebrafish embryogenesis. Values were normalized to the first time point (egg). Data is presented as mean with standard deviation. (C) Protein expression profiles of respiratory chain subunits during zebrafish embryogenesis detected by shot-gun mass-spectrometry in isolated mitochondria. (D-G) Complex I, II, IV and V in-gel activity assay for isolated mitochondria from zebrafish embryos at different developmental stages. Left panel: Activity measurements in BNP. Purple (D and E), brown (F) or black (G) staining indicates complex activity. Right panel: Quantification of complex activity normalized to complex abundance. Complex abundance was assessed by Coomassie staining.

### Large-scale changes to the mitochondrial proteome and phospho-proteome during embryogenesis

Having ruled out limited NADH availability and changes in ETC complex levels, accessory factors or modifications of ETC subunits as mechanisms underlying the increase in ETC activity during embryogenesis, we assessed changes to the mitochondrial proteome and phospho-protein by isolating mitochondria at different time-points and performing shotgun proteomics and phospho-proteomics combined with TMT (**Figure 4** and **Supplementary Figure S3**). Overall, we confidently detected 1761 proteins in the label-free proteome measurements of isolated mitochondria (detected with at least 2 peptides), of which 709 are annotated as mitochondrial while 1052 are co-purified proteins, including ER, Golgi and cytoplasmic proteins. PCA analysis of the top 500 differentially expressed mitochondrial proteins (DEPs) revealed separation of samples by embryonic time (**Figure 4B**). In total, 954 proteins were differentially expressed (adjusted p-value < 0,05 and absolute log2-fold-change > 1) (**Figure 4C**). Differential enrichment analysis and k-means clustering identified five clusters of proteins exhibiting distinct expression profiles (**Figure 4C**). To determine whether DEPs shared common pathways or biological functions, we carried out Gene Ontology (GO) enrichment analyses. Three clusters stood out particularly: cluster 3 containing downregulated DEPs at 24 hpf was enriched for proteins associated with mitochondrial morphology. Examples of these proteins are the Drp1 adaptor Fis1^44–46^ and the outer membrane protein Slc25a46^47,48^, which had previously been implicated in regulating mitochondrial fission (**Figure S4A)**. This cluster also included mitochondrial ETC assembly factors and complex I subunits (**Supplementary Table S1, S2**), which is consistent with our BNP results (**Figure 3A, B**). On the other hand, cluster 4 and cluster 5 containing upregulated DEPs were enriched for cytoplasmic translation and endoplasmic reticulum proteins, respectively. This was particularly interesting since analysis of the total embryonic proteome revealed constant levels of cytoplasmic ribosomal subunits and ER proteins up to 24 hpf^37^ (**Figure 4E, F**), implying that these proteins become more associated with mitochondria during embryogenesis.

**Figure 4.**
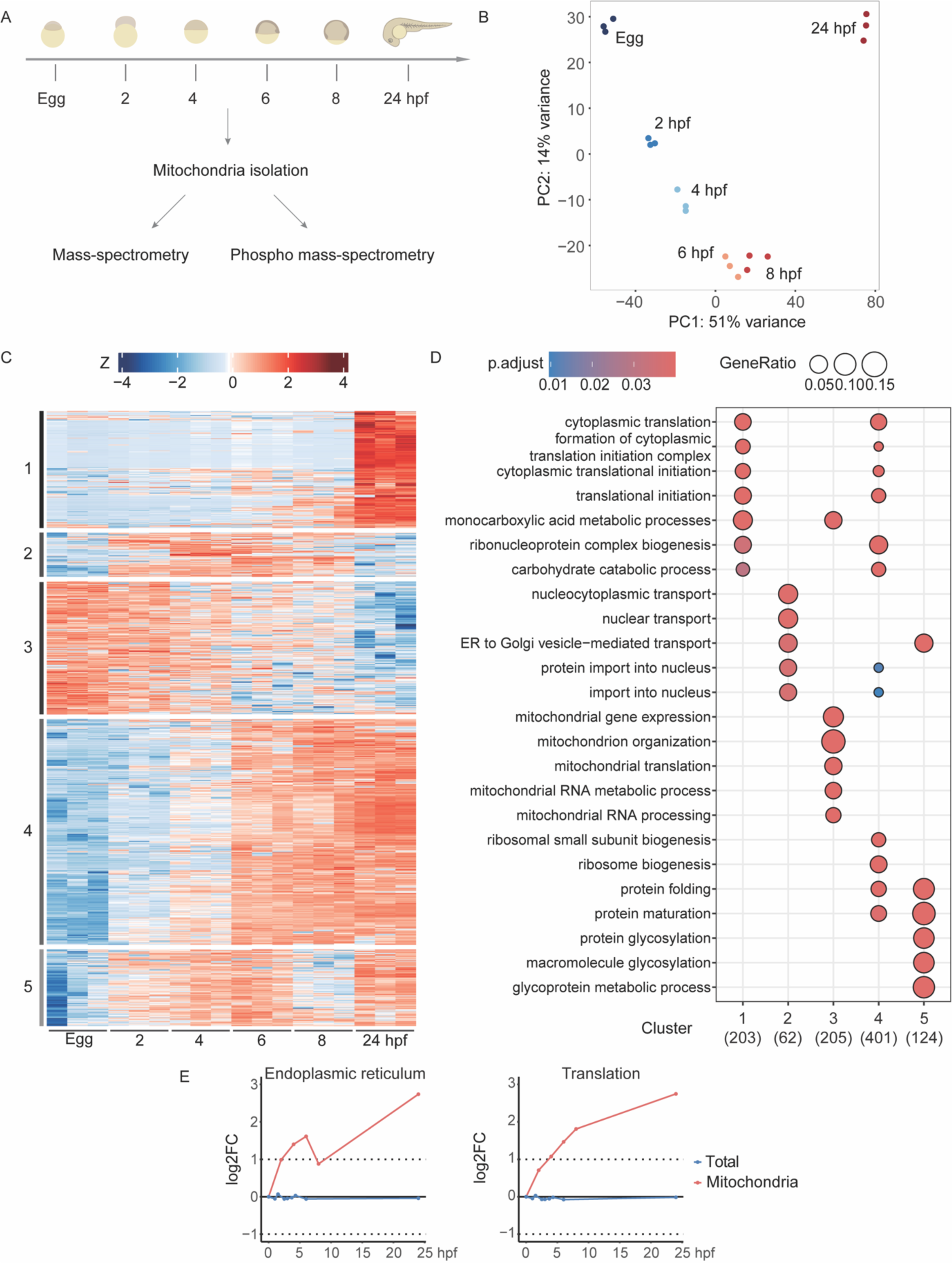
The mitochondrial proteome undergoes major changes during zebrafish embryogenesis. (A) Scheme of the experiment. (B) PCA plot for the top 500 differentially expressed mitochondrial proteins. (C) Proteomics of zebrafish mitochondria isolated at different stages of embryogenesis. Heatmap of the subset of differentially regulated mitochondrial proteins. Proteins were classified as differentially regulated if their absolute log2FC > 1 and adjusted P-value < 0.05. The color code represents z-scores for each protein. Each row corresponds to an individual protein and one square is a single replicate (3 replicates for each stage). (D) Gene ontology enrichment analysis for the respective cluster. (E) Line plots of log2FC of mean protein expression normalized to the first time point of differentially expressed proteins belonging to two GO terms, ER (left) and translation (right), in whole embryos (total embryonic proteome) and isolated mitochondria during zebrafish embryogenesis.

PCA analysis of the top 500 differentially expressed phospho-peptides in isolated mitochondria revealed that the 24 hpf time-point was most distinct (**Figure S3A)**. Analysis of differentially phosphorylated peptides in mitochondria during embryogenesis revealed four clusters of proteins with different patterns of phosphorylation. In line with our mitochondrial proteomics findings, proteins subject to differential phosphorylation during development included key factors related to mitochondrial fission, such as Drp1 and its adaptors Mff, and Mief1^49^, as well as Slc25a46 (**Supplementary Table S3**). Notably, while our mitochondrial proteome revealed that the total protein amount of Drp1 increases during zebrafish embryogenesis, Drp1 phosphorylation was reduced at later time points (**Figure S4B**) at the conserved position S616^50^, which is known to facilitate mitochondrial fission^50,51^. Our comprehensive analysis of mitochondrial proteomics and phospho-proteomics, coupled with GO term enrichment analysis, raised the interesting possibility that changes in mitochondrial morphology and association between endoplasmic reticulum and mitochondria may contribute to the observed increase in mitochondrial activity during embryogenesis.

### Mitochondria elongate during later stages of embryogenesis

It has previously been observed that elongated mitochondria have a higher oxygen consumption rate than fragmented mitochondria^52–54^. Interestingly, our proteomics data indicated a decrease in the abundance of mitochondrial morphology factors, including Drp1 adaptor, together with a decrease in the phosphorylated form of Drp1 (**Figure S4B**). This suggests that mitochondria might also elongate during zebrafish development, similar to what has been observed in other organisms^14–16^. We therefore analyzed mitochondrial morphological changes in zebrafish embryos at high spatio-temporal resolution.

To this end, we generated transgenic zebrafish lines with fluorescently labelled mitochondria (mitochondrial targeting sequence mts-EGFP or mts-DsRed) to visualize mitochondria by time-lapse spinning disk confocal imaging from 6 to 21 hpf.

We observed drastic morphological changes: Mitochondria were initially round and fragmented but started to elongate during somitogenesis (10-12 hpf) (**Figure 5A, movie 1**). To quantify changes in mitochondrial morphology over time, we used MitoSegNet^55^, a deep learning model for quantifying mitochondrial morphology that we trained on our dataset (**Figure S4C**). Despite embryo-to-embryo differences in the extent of elongation, we consistently detected an increase in the average size of mitochondria and decreases in mitochondrial number and circularity during embryogenesis (**Figure 5B-D**). To examine mitochondrial morphology during earlier developmental stages, we fixed zebrafish eggs and embryos at 2, 4 and 6 hpf, which revealed fragmented mitochondria at all stages, consistent with our live imaging analyses starting during gastrulation (**Figure S4D**). Additionally, in activated eggs and 16-cell stage embryos (2 hpf), mitochondria were found to cluster together (**Figure S4D**).

**Figure 5.**
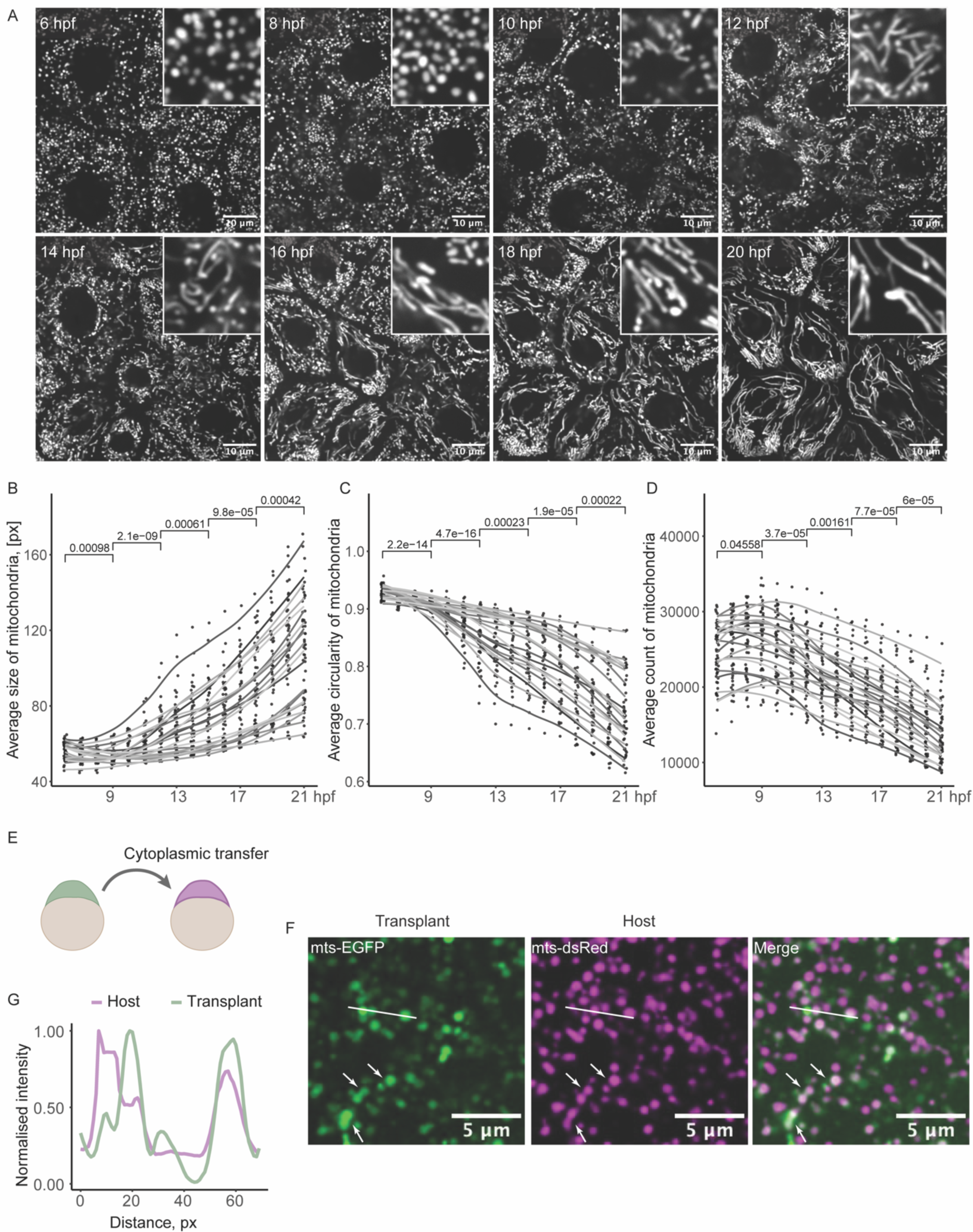
Fragmented yet fusion-competent mitochondria elongate during zebrafish embryogenesis. (A) Representative images of spinning disk confocal time-lapse microscopy of mts-dsRed labeled mitochondria during zebrafish embryogenesis starting from 6 hpf. The first mitochondrial morphological changes are observed around 10-12 hpf. Scale bar corresponds to 10 µm. (B-D) Morphological quantification of mitochondria using the MitoSegNet segmentation tool (**Figure S4A**). Average size (B), circularity (1 corresponds to a perfect sphere) (C) and number of mitochondria (D) were measured in segmented images of zebrafish mitochondria at different developmental stages. Each dot represents the average measurement within an image slice; 3 slices were imaged per embryo. Grey lines represent the nonparametric regression for each embryo. (E) Scheme of the transplantation experiment. Cytoplasm containing mts-EGFP labeled mitochondria was transferred to the embryo with mts-dsRed labeled mitochondria. (F) Representative images of the transplantation experiment, showing the overlap of mts-dsRed and mts-EGFP labeled mitochondria. Scale bar corresponds to 5 µm. (G) Intensity profiles of host (magenta) and transplanted (green) mitochondria along the white line in (F).

The fragmented mitochondria observed in the early stages could indicate either a lack of mitochondrial fusion or predominance of fission over fusion during these early stages. To distinguish between these two possibilities, we determined whether mitochondria could fuse at early stages by transferring cytoplasm containing green-labelled mitochondria into a stage-matched 1-hour embryo with red-labelled mitochondria (**Figure 5E-F**). One hour after transplantation we detected mitochondria labelled with both fluorophores (**Figure 5D, E**), supporting the idea that mitochondria generally can fuse in early embryos. However, this fusion activity appeared to be consistently counterbalanced by higher fission activity, as evidenced by increased levels of the active form of Drp1, phosphorylated at S616, in early embryos (**Figure S4B**).

Our findings showed substantial alterations in mitochondrial morphology, with predominantly fragmented yet fusion-competent mitochondria in the early embryo that start to elongate during somitogenesis, which correlates with the increase in OCR during later stages.

### Increase in ER-mitochondria association during early stages of embryogenesis

Mitochondrial morphology changes only started during somitogenesis and therefore could not account for the rise in mitochondrial activity during early stages (**Figure 1B**), suggesting the presence of additional mechanism(s) contributing to the early increase in mitochondrial activity. Interestingly, in our mitochondrial proteomic analysis we observed an early increase (cluster 5) in the levels of endoplasmic reticulum (ER) proteins during embryonic development (**Figure 4C**) in the absence of a concomitant increase of ER protein levels in the total proteome (**Figure 4F**), suggesting a potential increase in the interaction between ER and mitochondria. The interaction between mitochondria and ER is known to regulate mitochondrial homeostasis by providing metabolites such as Ca^2+^ and glycerophospholipids^56–59^, which could contribute to the increase in mitochondrial activity. Consistent with an increased ER-mitochondrial interaction, analyses of our proteomics data of isolated mitochondria revealed an increase in proteins such as Vapb^60,61^ and Mospd1^62^, which are known to mediate mitochondria-ER interaction (**Figure S5A**). Moreover, we detected differential phosphorylation of ER proteins important for Ca2+ transport between the ER and mitochondria, such as Pdzd8^63^, Fkbp8^64^ and Canx^65^, during development (**Figure S5B**), and upregulation of the essential mitochondrial calcium uniporter (MCU) regulator Smdt1b^66^ right after fertilization (**Figure S5A**).

To assess the relative distribution of mitochondria and ER during early embryogenesis, we generated transgenic fish lines with both ER and mitochondria fluorescently labelled. Time-lapse imaging of dual-labelled embryos revealed that mitochondria and ER were largely segregated into different compartments during the cleavage divisions, with mitochondria preferentially localizing to the cell periphery and ER being present more in the center of each cell (**Figure 6A**, **movie 2**).

**Figure 6.**
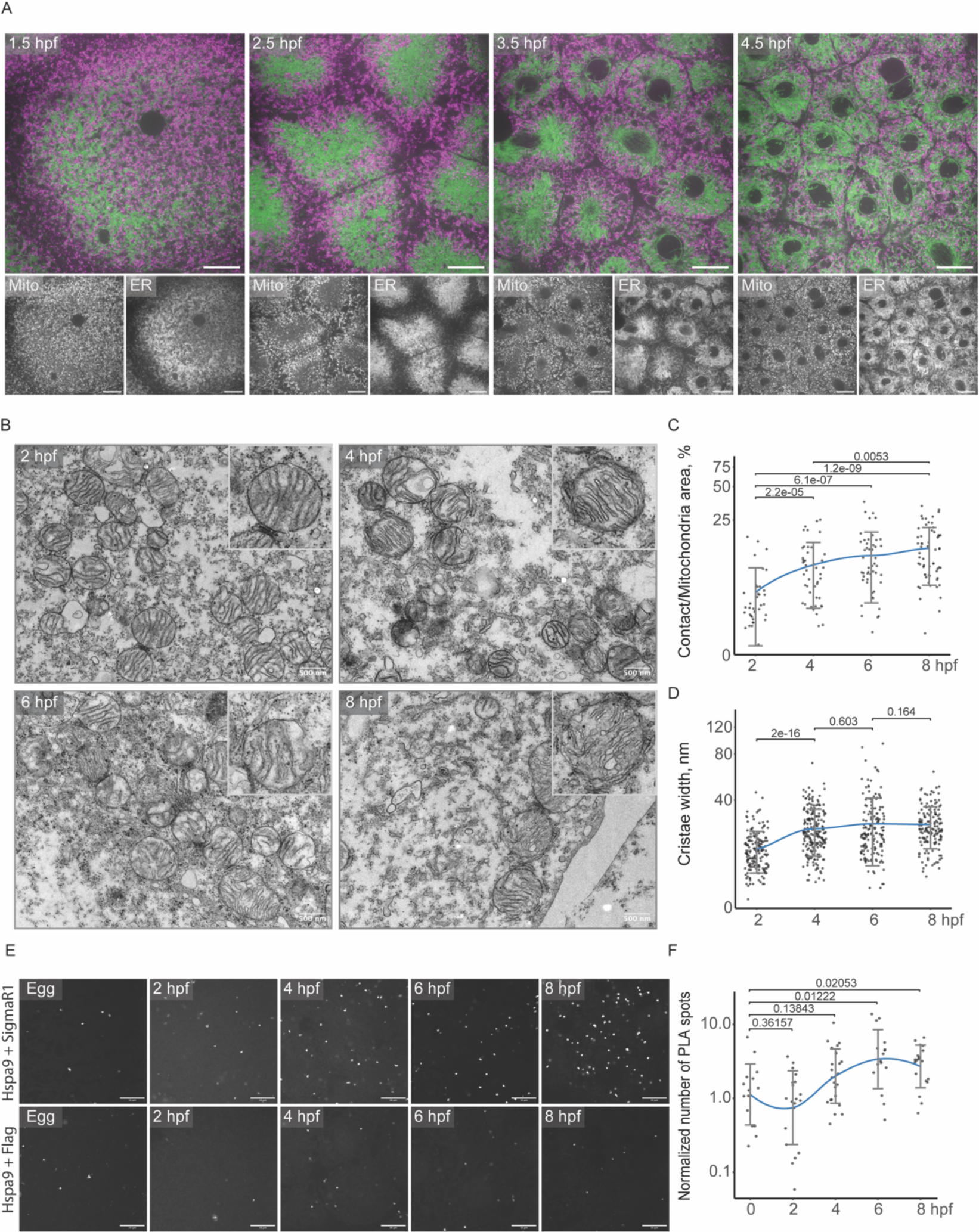
Mitochondria-ER association increases during early stages of zebrafish embryogenesis. (A) Representative images of spinning disk confocal time-lapse microscopy of dual-labelled embryos (mts-dsRed labeled mitochondria (magenta); EGFP labeled ER (green)) during zebrafish embryogenesis starting from 1.5 hpf. Scale bar corresponds to 20 µm. (B) Representative electron microscopy images of zebrafish embryos at different developmental stages. Scale bars correspond to 500 nm. (C) Quantification of the ER-mitochondria contact area during early zebrafish embryogenesis (the y-axis is scaled logarithmically). The blue line represents the nonparametric regression with standard deviation. Kruskal-Wallis one-way ANOVA P-values (Benjamini-Hochberg correction) are indicated for statistically significant changes. (D) Quantification of mitochondrial cristae width during early zebrafish embryogenesis (the y-axis is scaled logarithmically). The blue line represents the nonparametric regression with standard deviation. Kruskal-Wallis one-way ANOVA P-values (Benjamini-Hochberg correction) are indicated for statistically significant changes. (E) Representative images of *in situ* PLA performed with antibody pairs detecting mitochondrial (anti-Hspa9) and ER (anti-SigmaR1) proteins as well as control conditions (anti-Hspa9 and anti-Flag) during early zebrafish embryogenesis. Scale bar corresponds to 10 µm. Only a small area of each embryo image is shown. Full embryo images are shown in **Figure S5C**. (F) Quantification of the number of PLA spots (the y-axis is scaled logarithmically). The blue line represents the nonparametric regression with standard deviation. Data was normalized by dividing the values of each time point by the mean value of the corresponding control time point (see **Figure S5C, D**). Kruskal-Wallis one-way ANOVA P-values (Benjamini-Hochberg correction) are indicated for statistically significant changes.

To assess potential changes in the association between mitochondria and ER during early embryogenesis, we used two independent assays to quantify the occurrence of ER-mitochondria interactions over time: transmission electron microscopy (TEM) and a proximity ligation assay (PLA). TEM-images of sections of embryos at early embryonic stages (2 hpf, 4 hpf, 6 hpf and 8 hpf) revealed an increase in the fraction of the mitochondrial surface covered by the ER during early development (**Figure 6B, C**). Additionally, the width of the mitochondrial cristae also increases (**Figure 6D**), which correlates with the rise in respiration during embryogenesis (**Figure 1B**) and has previously been linked to an increase in ETC activity and ATP production^67^.

The PLA also supports an increase in ER-mitochondria association during early embryogenesis. When applying antibodies against SigmaR1 (ER resident) and Hspa9 (outer mitochondrial membrane protein), the number of foci increased, indicating rising proximity of the organelles from 4 hpf on (**Figure 6E, F** and **Figure S5A**). While some background signal was also detected using the control anti-Flag antibody, the number of spots increased over time only in the presence of both the ER- and mitochondrial-specific antibodies (**Figure 6E** and **S5A, B**) as evident after normalization of the Hspa9–SigmaR1 to the Hspa9–Flag (background) derived signal (**Figure 6F**). The observed increase in the number of spots in the PLA assay is consistent with an increased proximity between mitochondria and ER at early stages of development.

Together, these results suggest an increase in ER-mitochondrial interactions during early embryogenesis, which correlates with the early increase in mitochondrial activation.

## Discussion

### The mechanisms contributing to the activation of mitochondria during embryogenesis have not been systematically investigated

The early embryo represents a non-conventional cellular state characterized by dormancy, where many major energy-consuming processes, such as nuclear transcription^68,69^ and cytoplasmic translation^70,71^, are quiescent or occur at very low levels. Thus, it is not surprising that mitochondrial activity is generally low in early embryos across various organisms and increases later during development^11,27,28^. Mitochondrial function during embryogenesis has been studied for several decades, and multiple mechanisms are known to regulate the activity of the respiratory chain^72^. However, a comprehensive analysis of the molecular and cellular mechanisms that contribute to the continuous increase in mitochondrial activity during embryogenesis had not been performed in any organism. To address these gaps in knowledge, we provide a systematic characterization of molecular and cellular changes during embryonic mitochondrial activation in zebrafish.

### Identification of two sequential mechanisms of mitochondrial activation: an early increase in mitochondrial-ER association and later mitochondrial elongation

In this study, we find that mitochondrial activity increases continuously during embryogenesis in zebrafish (**Figure 1B**) and that both mitochondrial membrane potential and mitochondrial ATP production are essential for developmental progression (**Figure 1D**). Activation of mitochondria during early embryonic stages occurs in the presence of constant mitochondrial DNA content (**Figure 1C**), availability of the ETC substrate NADH (**Figure 2**), and in the absence of an increase in the abundance or enzymatic activity of ETC complexes (**Figure 3**). These findings imply that the rise in mitochondrial activity during embryogenesis relies on mechanisms that differ from the previously described ones during oogenesis in mice^12^ and flies^13^, where the abundance of respiratory chain complexes was shown to be downregulated to reduce mitochondrial activity in oocytes.

Indeed, changes to the mitochondrial proteome (**Figure 4**) followed up by detailed morphological analyses pointed towards two mechanisms as possible contributors to the increase in mitochondrial activity during embryogenesis (**Figure 4E**): an increase in mitochondria-ER association during early stages (**Figure 6**) and dynamic changes to the mitochondrial morphology, leading to their fusion and elongated shape during later stages (**Figure 5**).

### Increased association between mitochondria and ER as a new mechanism regulating mitochondrial activity

Our mitochondrial proteomics data revealed the increase in abundance of ER proteins, starting as early as 2 hours post fertilization. This finding was particularly interesting since ER proteins showed constant levels in proteomic experiments from total embryos^37^ (**Figure 4E**), indicating increased proximity and interaction between mitochondria and the ER over time and thus increased co-purification of ER proteins with mitochondria.

Reorganization of both the ER and mitochondria have been observed and associated with oocyte maturation. During murine oogenesis, the cytoplasmic ER network observed in germinal vesicle-stage oocytes (GV) has been described to transform into distinct cortical clusters in metaphase II eggs (MII)^73,74^. These clusters are thought to be essential for oocyte activation by facilitating the generation of Ca^2+^ transients^75^. Simultaneously, mitochondria in mice were shown to organize into mitochondria-associated ribonucleoprotein domains (MARDOs) in the cytoplasm, which are depleted of ER and formed to store maternal mRNAs^76^. This reorganization suggests that mitochondrial and ER interactions may be reduced during oogenesis. Upon egg activation and during the first cleavage divisions, the rearrangements of these organelles observed by imaging fluorescently labeled ER and mitochondria (**movie 2**) may affect mitochondria-ER interaction. In support of this notion, mitochondria in activated oocytes and early embryos were found in clusters, which gradually dispersed as development progressed (**Figure S4D**).

The interaction between the ER and mitochondria is important for mitochondrial homeostasis as the ER is an intracellular storage site for Ca^2+^, which is utilized by mitochondria to regulate essential functions, including metabolism and energy production. In addition, Ca^2+^ is known to directly activate some Krebs cycle enzymes as well as complex V^21,77,78^. For example, pyruvate dehydrogenase phosphatase^79^, which removes an inhibitory phosphate from the E1A subunit of pyruvate dehydrogenase, depends on Ca^2+^ as do isocitrate dehydrogenase and oxoglutarate dehydrogenase, which convert isocitrate to α-ketoglutarate and α-ketoglutarate to succinyl-CoA, respectively^77,78^. The increase in the levels of citrate, aconitic acid and α-ketoglutarate as detected in our metabolic profiling during embryogenesis correlates with the increase in ER-mitochondria interaction, which is in line with enhanced ER-mitochondrial interaction activating mitochondria.

Additional support comes from our observation that ER proteins mediating Ca^2+^ transfer between ER and mitochondria are differentially phosphorylated during embryogenesis (**Figure S5A, B**). While further experiments will be required to directly test a functional link between the increased ER-mitochondria interaction and the observed changes in phosphorylation of ER proteins as well as more generally the increase in mitochondrial activity, our data provide a strong basis for this previously largely unexplored feature.

### What drives the increase in mitochondrial fusion, and to which extent does it contribute to the increase in mitochondrial activity during embryogenesis?

Another major change detected in our mitochondrial proteomics data was the downregulation of proteins linked to mitochondrial organization and morphology (**Figure 4C**). In line with this, we found that mitochondria undergo large-scale changes to their morphology, leading to mitochondrial fusion and elongation at later stages of embryogenesis (**Figure 5A-D**). Closer inspection of the dynamics of differentially expressed proteins implicated in mitochondrial morphological changes showed that several proteins linked to mitochondrial fission, such as the adaptor of the known fission factor Drp1, Fis1 and the outer membrane protein Slc25a46, were downregulated during embryogenesis, consistent with the observed mitochondrial elongation and fusion over time. However, the increased protein abundance of Drp1, and the decreased abundances of the known fusion factors Mfn2, Mfn1b and Opa1, were counter-intuitive at first sight (**Figure S4A**). Two observations could explain these seemingly contradictory findings: First, our transplantation experiment showed that mitochondria also fuse in early embryos (**Figure 4E-G**). Second, and in line with our morphological analyses that fission is more active than fusion in the early embryo, we found that the (activating) S616-phosphorylated form of Drp1 is more abundant during early embryonic stages than its non-phosphorylated form (**Figure S4B**). These data support the idea that the fission/fusion balance is shifted towards fission in the early embryo, which keeps them fragmented up to 10-12 hpf.

Elongated mitochondria have been reported to exhibit higher rates of respiration in many instances^52–54^. It would therefore be interesting to investigate in future studies to which extent mitochondrial elongation contributes to the increase in mitochondrial activity during development. Possible avenues to explore are modulating Drp1 level and/or activity identifying the phosphatase responsible for dephosphorylating S616 of Drp1 during development. Additionally, there may be other mechanisms regulating Drp1 activity; for instance, it has been reported that the membrane composition of the outer mitochondrial membrane can influence Drp1 activity^80^. For example, cardiolipin was shown to stimulate the GTPase activity of Drp1, thereby promoting mitochondrial fission^80^. Interestingly, our mitochondrial proteomics data indicates that Plscr3b^81^, which has been implicated in facilitating the transport of cardiolipin from the inner to the outer mitochondrial membrane, is upregulated during embryogenesis, making it an additional candidate for future functional studies.

Overall, in light of the fact that mitochondrial fusion appears to be a conserved feature of the dynamic changes to the mitochondrial morphology during embryogenesis across multiple organisms^14–16^, we can speculate that also the increase in mitochondrial-ER interaction may be a universal feature that contributes to the increase in mitochondrial activity during embryogenesis.

## Acknowledgements

We thank Karin Panser, Carina Pribitzer and Lena Bohaumilitzky for experimental support; Jodi Nunnari, Thomas Hurd, Antonio Zorzano, Lenka Radonova, Carmen Williams, Styliani Panagiotou, Olof Idevall, Anna Korte, Efe Ilker and Jonathan Rodenfels for fruitful discussions, feedback and providing reagents; the Proteomics Facility at IMP/IMBA/GMI using the VBCF instrument pool, and K. Stejskal and S. Opravil for the processing of MS samples; the BioOptics facility under the lead of Karin Aumayr for their help with microscopy data acquisition and analysis; the Metabolomics and Electron Microscopy Facilities at Vienna BioCenter Core Facilities (VBCF), member of the Vienna BioCenter (VBC), for support in metabolomics and EM imaging data acquisition; the IMP animal facility, especially F. Puhl, K. Rattner, J. König and D. Sunjic for their care of zebrafish; T. Mylenko, A. Bandura, G. Deneke, and M. Binner for their help with genotyping; the Molecular Biology Service at IMP for Sanger sequencing and for providing competent cells and reagents; and the entire Pauli lab for critical feedback and valuable discussions. Work in the Pauli lab was supported by funding from the IMP, which receives institutional funding from Boehringer Ingelheim and the Austrian Life Sciences Program 2023 (# 48924910), as well as the FWF START program (Y 1031-B28), the ERC CoG (101044495/GaMe), the HFSP Career Development Award (CDA00066/2015), a HFSP Young Investigator Award (RGY0079/2020) and the FWF SFB RNA-Deco (project number F80). AC was supported by an MSCA-IF-EF-SE (895790) and by the European Union’s Framework Programme for Research and Innovation Horizon 2020 (2014-2020) under the Marie Curie Skłodowska Grant Agreement Nr. 847548. BA was supported by the Austrian Science Fund (FWF) DK W1212 and A.B. was supported by an ERC Starting Grants (#677006). For the purpose of Open Access, the authors have applied a CC BY public copyright license to any Author Accepted Manuscript version arising from this submission.

## Author contributions

**Anastasia Chugunova**: conceptualization, data curation, formal analysis, funding acquisition, investigation, methodology, project administration, supervision, validation, visualization, writing – original draft. **Hannah Keresztes, Roksolana Kobylinska and Marcus Strobl**: data curation, investigation, methodology. **Maria Novatchkova**: formal analysis, software (computational analyses). **Thomas Lendl**: software (MitoSegNet). **Michael Schutzbier**, **Gerhard Dürnberger, Richard Imre, Elisabeth Roitinger**: data curation, formal analysis, resources, supervision (mass-spectrometry). **Pawel Pasierbek, Alberto Moreno Cencerrado**: resources, supervision (live-imaging)**. Marlene Brandstetter**: data curation, formal analysis, resources (electron-microscopy). **Thomas Köcher**: data curation, formal analysis, resources (metabolomics). **Benedikt Agerer**, **Jakob-Wendelin Genger**, **Andreas Bergthaler:** data curation, formal analysis, resources (Seahorse measurements). **Andrea Pauli**: conceptualization, data curation, funding acquisition, project administration, resources, supervision, writing.

## Supplementary Figures

**Figure S1.**
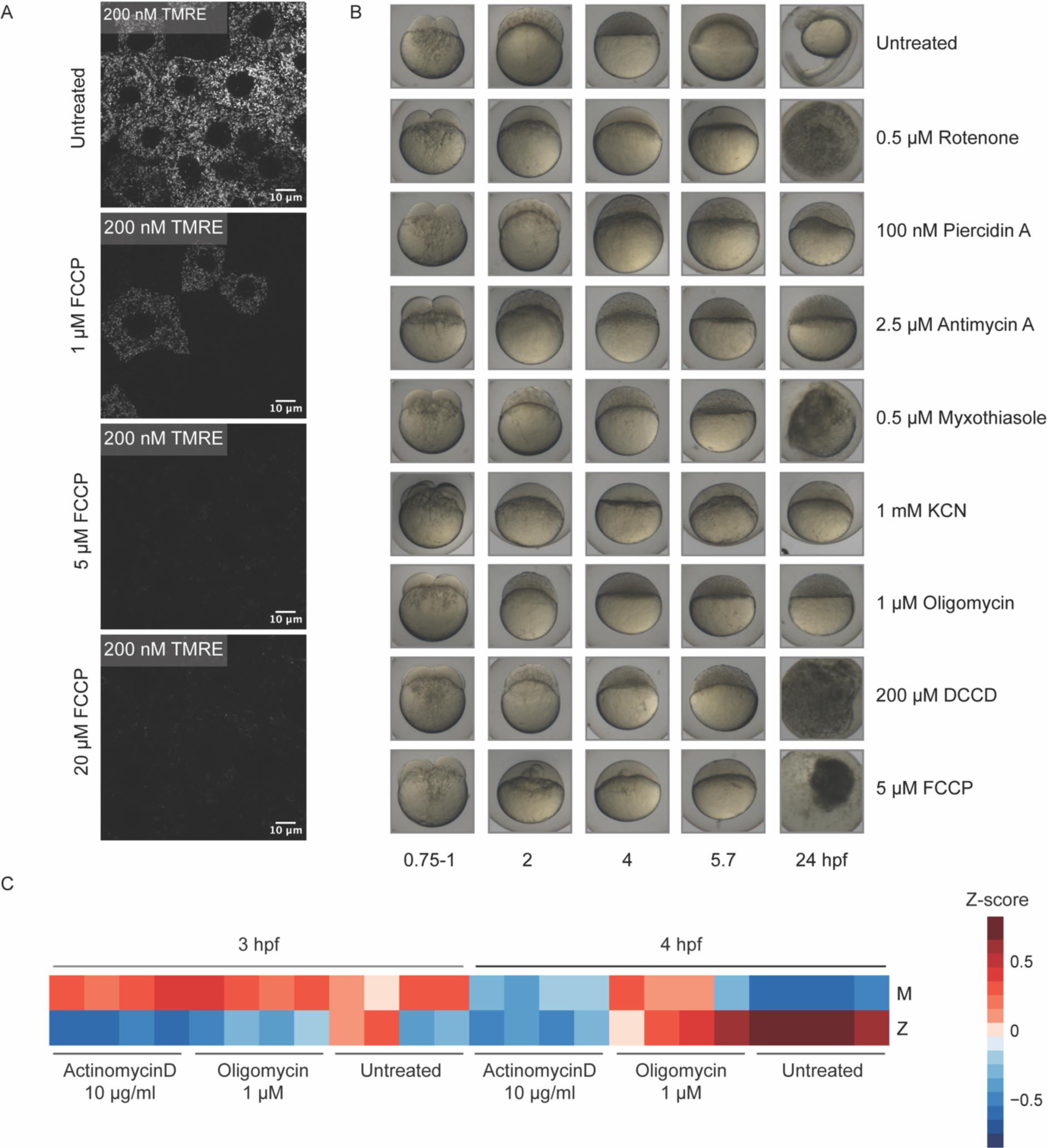
Zebrafish embryo development is affected by treatment with different respiratory chain inhibitors. (A) Effect of different concentrations of FCCP (0-20 µM) on embryos injected with 200 nM TMRE. TMRE was used as an indicator of the presence of mitochondrial membrane potential. A concentration of 5 µM FCCP was found to block mitochondrial membrane polarization. (B) Effects of respiratory chain inhibitors on zebrafish embryo development. (C) Average RNA expression of maternal (M) and zygotic (Z) genes in wild-type embryos, in embryos treated with Oligomycin (inhibition of ATP production by inhibiting complex V), and in embryos injected with actinomycin-D (inhibition of transcription) at 3 and 4 hpf. The color code represents z-scores. Each square corresponds to a single replicate (4 replicates for each treatment and stage).

**Figure S2.**
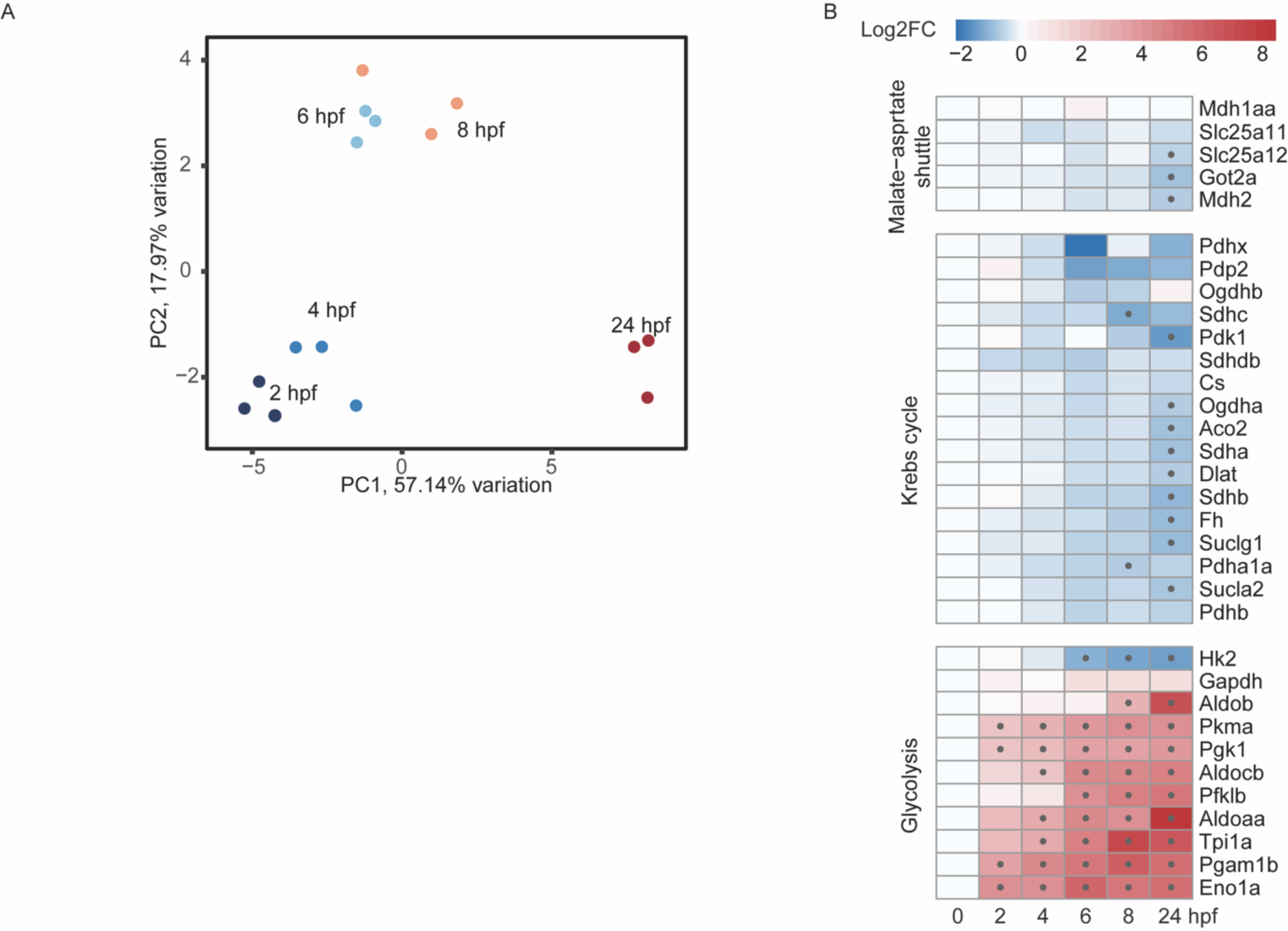
Metabolomics of zebrafish embryos during embryogenesis. (A) PCA plot for detected metabolites at different stages of embryogenesis. (B) Protein expression profile of malate-aspartate shuttle subunits, Krebs cycle enzymes and glycolysis enzymes during zebrafish embryogenesis in isolated mitochondria. The color code represents log2FC relative to the first time point 0 (Egg). The dot represents significant values (with adjusted P-value < 0.05 and absolute log2FC > 0.5).

**Figure S3.**
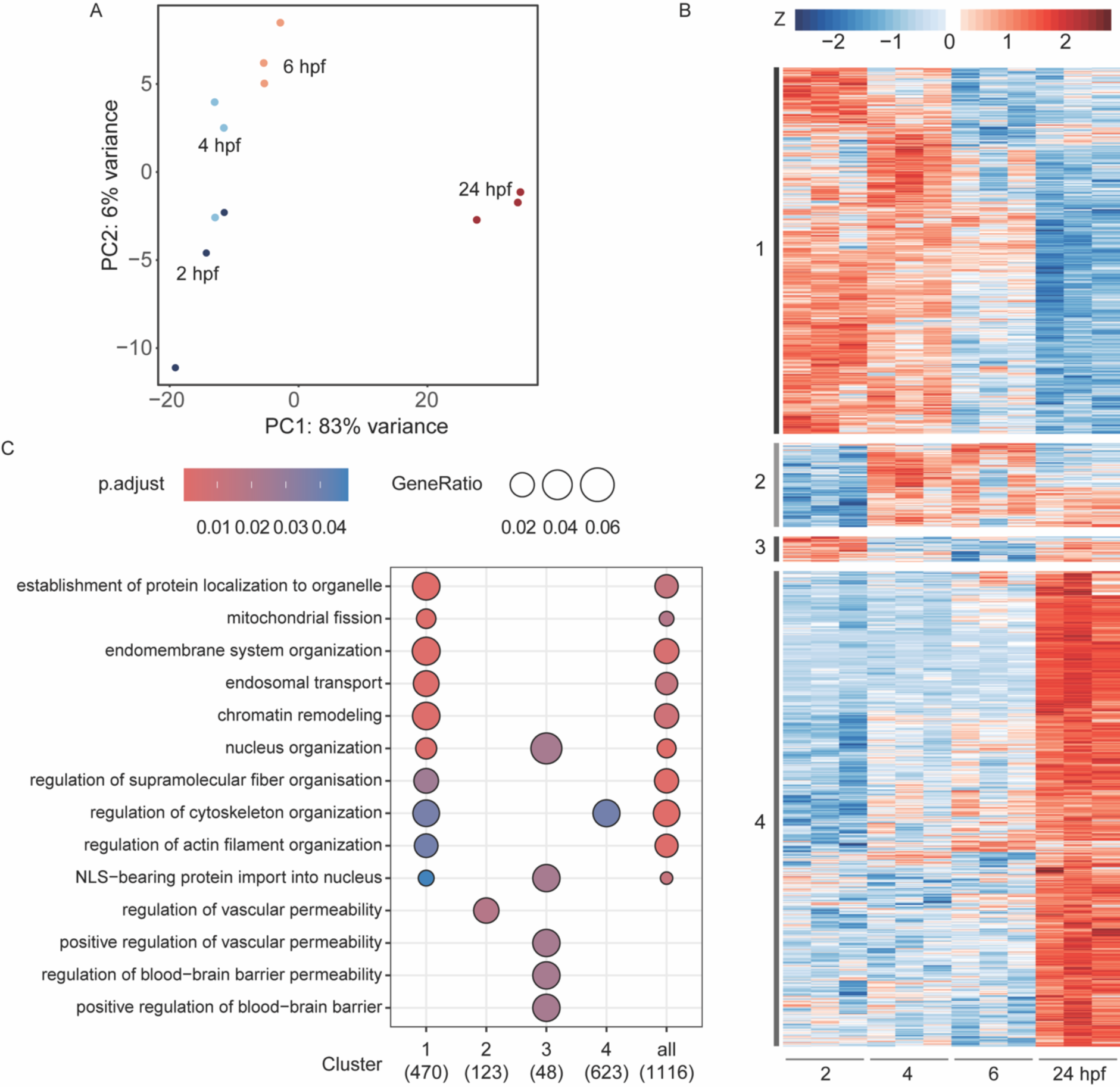
Mitochondrial proteomics and phospho-proteomics during zebrafish embryogenesis. (A) PCA plot for the top 500 differentially phosphorylated peptides. (B) Phospho-proteomics of zebrafish mitochondria isolated at different stages of embryogenesis. Heatmap of the subset of differentially phosphorylated peptides normalized to the abundance of the respective protein during embryogenesis. Phospho-peptides are classified as differentially regulated if their adjusted P-value < 0.05. The color code represents z-scores for each protein. Each row corresponds to an individual phospho-peptide and one square is a single replicate (3 replicates per stage). (C) Gene Ontology enrichment analysis for each cluster.

**Figure S4.**
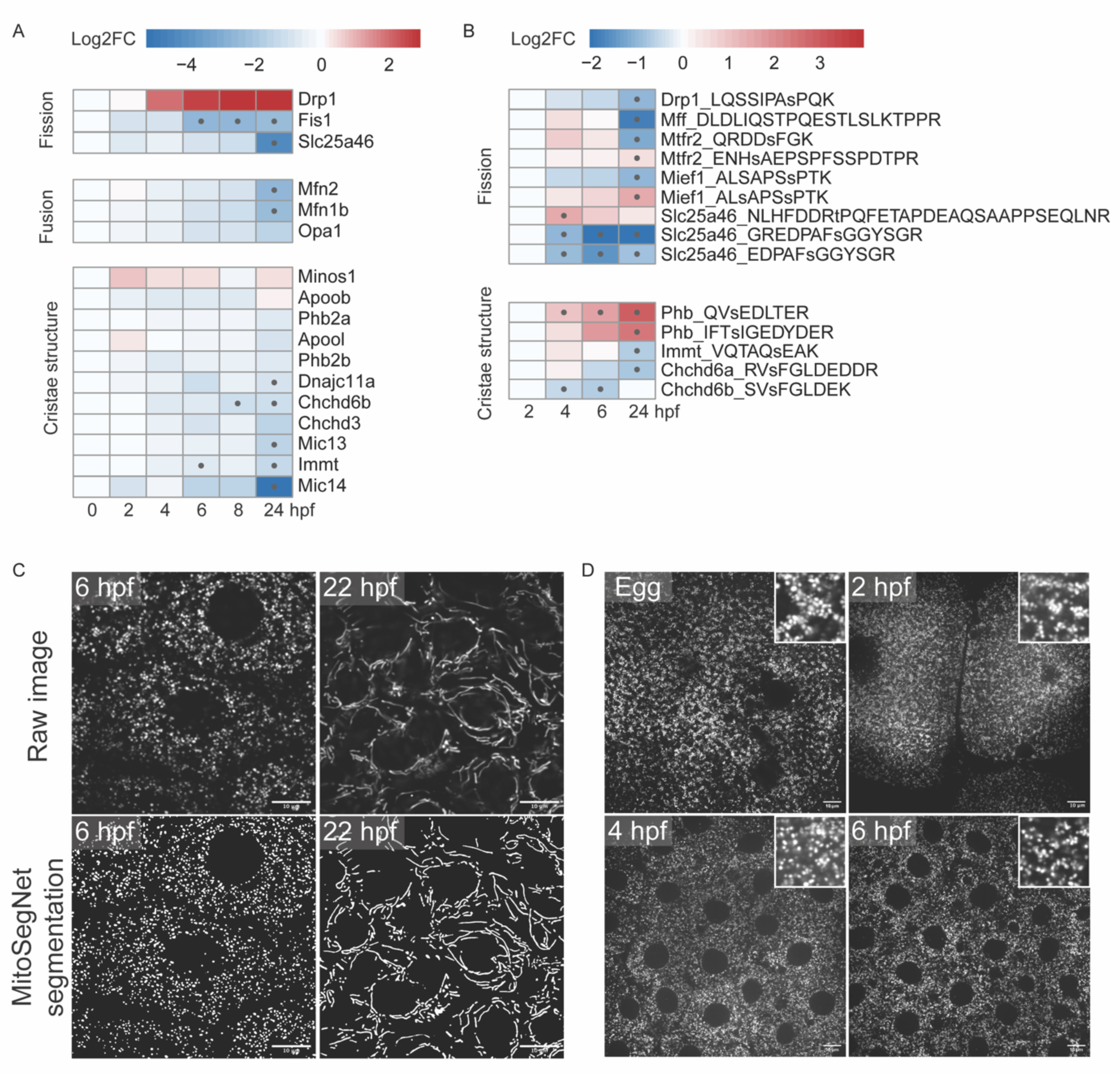
Analysis of mitochondrial morphological changes during zebrafish embryogenesis. (A) Protein expression profiles of proteins associated with fission, fusion, and cristae structure during zebrafish embryogenesis in isolated mitochondria. The color code represents log2FC relative to the first time point 0 hpf (Egg). The dot represents significant values (with adjusted P-value < 0.05 and absolute log2FC > 0.5). (B) Relative amounts of phospho-peptides normalized to the abundance of the corresponding proteins for a subset of the proteins associated with fission and cristae structure. The color code represents log2FC relative to the first time point 2 hpf. The dot represents significant values (with adjusted P-value < 0.05). (C) Representative images of mitochondrial segmentation with the MitoSegNet^55^ tool. Raw images are shown at the top, and MitoSegNet segmentations at the bottom. The scale bar is 10 µm. (D) Representative images of mts-EGFP labelled mitochondria in the fixed zebrafish eggs and in fixed embryos at early stages of embryogenesis. Scale bar corresponds to 10 µm.

**Figure S5.**
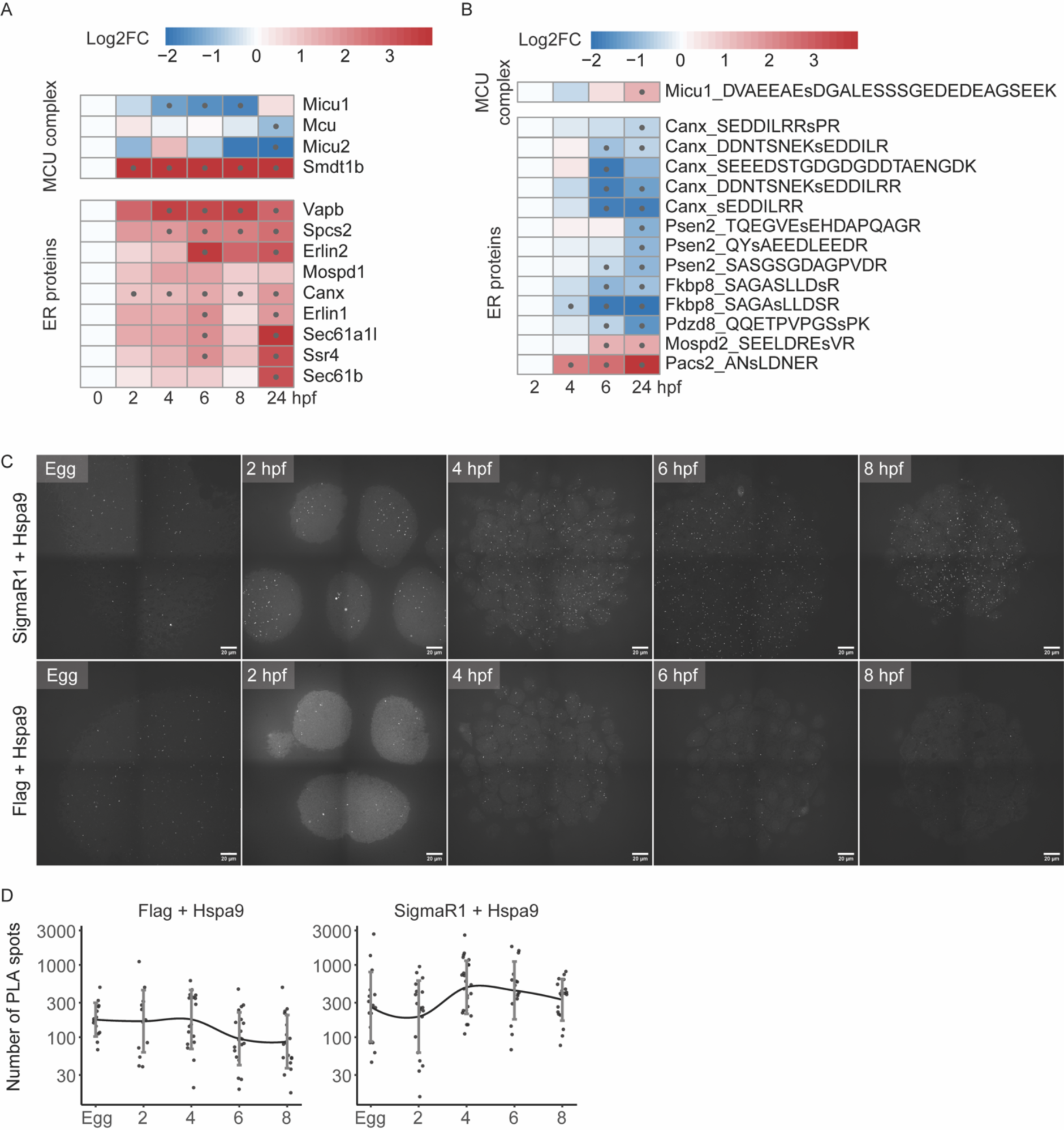
Mitochondria-ER proximity increases during embryogenesis. (A) Protein expression profiles of proteins associated with the MCU complex and ER during zebrafish embryogenesis in isolated mitochondria. The color code represents log2FC relative to the first time point 0 hpf (Egg). The dot represents significant values (with adjusted P-value < 0.05 and absolute log2FC > 0.5). (B) Relative amounts of phospho-peptides normalized to the abundance of the corresponding proteins for a subset of the proteins associated with the MCU complex and ER. The color code represents log2FC relative to the first time point 2 hpf. The dot represents significant values (with adjusted P-value < 0.05). (C) Representative images of *in situ* PLA performed with the antibody pairs anti-Hspa9 (mitochondrial protein) and anti-SigmaR1 (ER protein) (top) and anti-Hspa9 and IgG control (bottom) during early zebrafish embryogenesis. Scale bar corresponds to 20 µm. (D) Quantification of the number of PLA spots for samples treated with antibody pairs anti-Hspa9 and IgG control (left) and anti-Hspa9 and anti-SigmaR1 (right) during early zebrafish embryogenesis. The blue line represents the nonparametric regression with standard deviation.

**Supplementary Table S1**

Label-free quantification of mitochondrial shot-gun proteomics (related to Figure 3, 4, S4, S5).

Column 1 – Description for proteins Column 2 – Protein name

Columns 3-20 – Relative amounts of phospho-peptides normalized to the abundance of the corresponding proteins (Egg, 2, 4, 6, 8 and 24 refer to hours post fertilization, and the _1/2/3 to the three biological replicates)

**Supplementary Table S2**

Differential protein expression analysis of mitochondrial shot-gun proteomics (related to Figure 3, 4, S4, S5).

Column 1 – Gene ID

Column 2 – Protein name

Column 3 – Protein ID

Columns 4-13 – log2FC and adjusted P-values

**Supplementary Table S3**

TMT quantification and differential peptide phosphorylation analysis of mitochondrial phospho-proteomics (related to Figure S3).

Column 1 – Protein ID

Column 2 – Peptide sequence with indications of modifications

Column 3 – Description for proteins

Column 4 – Description of modifications

Columns 5-16 - Relative amounts of phospho-peptides normalized to the abundance of the corresponding proteins (2, 4, 6, and 24 refer to hours post fertilization, and the _1/2/3 to the three biological replicates)

Columns 17-22 – log2FC and adjusted P-values

**Movie 1**

Changes in mitochondrial morphology observed during zebrafish embryogenesis using spinning disk confocal microscopy. Scale bar corresponds to 10 µm. Time is shown in hours post fertilization.

**Movie 2**

Mitochondrial (magenta) and ER (green) localization changes observed during zebrafish embryogenesis by spinning disk confocal microscopy. Scale bar corresponds to 20 µm. Time is shown in hours post fertilization.

## Materials and Methods

### Zebrafish husbandry

Zebrafish (*Danio rerio*) were raised at 28° C with a 14/10 hour light/dark cycle. Wild-type TLAB fish correspond to fish obtained by crossing AB with the natural variant TL (TupfelLongfin). All fish experiments were conducted according to Austrian and European guidelines for animal research and approved by local Austrian authorities (protocols for work with zebrafish GZ342445/2016/12 and MA 58-221180-2021-16).

### Plasmid construction

To generate the plasmid with EGFP or DsRed targeted to the mitochondrial matrix, the mitochondrial targeting sequences of Atp5f1b (ATGTTGGGAGCTGTGGGACGCTGCTGCACTGGGGCTTTGCAGGCTCTCAAGCCCGGGGTTCACCCCCTGAAGGCTCTCAACGGAGCTCCATCTCTGTTTTCACGCAGGGATTATGCTGCTCCTGCCGCTGCTGCTGCCGCCGCCAGC) and the EGFP/DsRed sequence were PCR-amplified from zebrafish cDNA and from a plasmid, respectively, and cloned by Gibson assembly into a vector containing Tol2-integrations sites in between the zebrafish *actb2* promoter, *actb2* 5’ UTR and the SV40 late polyadenylation signal. To generate the plasmid with EGFP or mScarlet3 targeted to endoplasmic reticulum, the fusion of EGFP/mScarlet3 with N-terminal ER targeting sequence of N-lysozyme (ATGCTGCTATCCGTGCCGTTGCTGCTCGGCCTCCTCGGCCTGGCCGTCGCCGACCGGTCGCACACC) and C-terminal KDEL signal (AGATCGTACAAGAAGGACGAGCTGTA) was amplified from a plasmid and cloned by Gibson assembly into a vector containing Tol2-integrations sites in between the zebrafish *actb2* promoter, *actb2* 5’ UTR and the SV40 late polyadenylation signal.

### Zebrafish transgenic lines

To generate transgenic lines, 15 pg of each plasmid was co-injected with 35 pg of *Tol2* mRNA into 1-cell embryos. Adult fish were crossed with wild-type fish, and GFP- and DsRed-positive embryos were raised to adulthood.

### Oxygen consumption measurement

Individual embryos were placed into each well (1 per well) of an Agilent XF96e Spheroid microplate containing 180 µl of E3 medium (5 mM NaCl, 0.17 mM KCl, 0.33 mM CaCl_2_, 0.33 mM MgSO_4_, 10^−5^% methylene blue). Basal respiration was recorded on the Seahorse XF96e at 28 °C. This was achieved by switching off temperature control. Measurement cycle was as follows (3:00 Mixing, 0:30 Waiting, 3:00 Measuring; 3x).

### Phenotyping after inhibitor treatment

Embryos were collected as soon as they were laid and put into E3 medium (5 mM NaCl, 0.17 mM KCl, 0.33 mM CaCl_2_, 0.33 mM MgSO_4_, 10^−5^% methylene blue) containing inhibitors. Embryos were kept in this medium for the duration of the experiment. For this experiment we used already established concentrations for inhibitors: 0,5 µM rotenone (Millipore, 557368), 100 nM piercidin A (Cayman, CAY-15379-1-1), 2,5 µM antimycin A (Sigma-Aldrich, A8674), 0,5 µM myxothiazol (Sigma-Aldrich, T5580), 1 mM KCN (Sigma-Aldrich, 60178), 1 µM oligomycin (Millipore, 495455), 200 µM DCCD (Millipore, 8029540250), 5 µM FCCP (Sigma-Aldrich, C2920). Untreated embryos were kept alongside as controls. Manual examination of defects was performed blindly at every 2 hours for the first 8 hpf and then at 24 hpf. Pictures were taken under a dissection microscope and a Blackfly S USB 3 camera (serial number 18255235) using the FlyCapture2 software.

### Total mtDNA quantification

We used a previously established protocol^35^ with minor changes. Samples (one embryo or activated oocyte per sample) were collected and immediately snap-frozen in liquid nitrogen. After thawing, samples were lysed for 4 h in 250 µl lysis buffer containing 10 mM Tris-HCl pH 8, 10 mM EDTA, 0,2% Triton X-100, 200 µg/ml Proteinase K, 200 mM NaCl 75 mM NaCl, 50 mM and 20 mM HEPES, supplemented with 0.4% SDS at 55 °C. After incubation at 98 °C to inactivate proteinase K, 210 µl isopropanol was added and samples were precipitated overnight at −20 °C. After centrifugation at 14.000 rcf for 10 min at 4 °C, DNA was washed with 500 µl 70% Ethanol and the pellet was dissolved in 10 µl TE buffer.

To generate the standard curve, the mitochondrial *mt-nd1* gene (NADH dehydrogenase 1, mitochondrial) was amplified using primers mt-nd1-cDNA-f CCACTTAATTAACCCCCTAGCC and mt-nd1-cDNA-r ATGTTTGTGGGGGTAGACCA and cloned into the pSC-A vector using the StrataClone PCR Cloning Kit. To quantify mtDNA, primers mt-nd1-F TCAGGAAGACACATGACTTCTACTTC and mt-nd1-R CAAATGGTCCTGCTGCATAC were used.

After DNA extraction, the mtDNA copy number was measured via qPCR using a standard curve. qPCR was carried out in a total reaction volume of 20 µL. Due to the small volumes of DNA, no input amounts of DNA were known (as nanodrop measurement were not possible), yet each sample contained 1 embryo as input. In all experiments, the concentrations of the reaction mixture were similar: primer concentrations were 1 µM, and the reaction contained 1x master-mix of the GoTaq qPCR kit (Promega). Reactions were carried out on the CFX96 Touch Real-Time PCR Detection System (Bio-Rad) in 96-wells plates. Thermocycling parameters were as follows: 2 min 50 °C, 2 min 95 °C, followed by 40 cycles of 15 s 95 °C (denaturation) and 1 min 60 °C (annealing and extension). For analyzing non-specific amplification, a dissociation (melting) curve was generated: 15 s 95 °C, 15 s 60 °C, 15 s 60 °C, cooling down to 4 °C. Amplification results were converted to absolute mtDNA copy numbers using a standard curve.

### Targeted metabolomics

Embryos were dechorionated and deyolked manually. Cell caps (10 per sample) were collected in a 1.5 ml tube, and the extraction buffer containing 20% water, 40% acetonitrile, 40% methanol was added. Metabolite extracts were analyzed either by reversed phase chromatography or by hydrophilic interaction liquid chromatography (HILIC), coupled to tandem mass spectrometry (LC-MS/MS), employing selected reaction monitoring (SRM). In HILIC, 1 µl of each sample was injected onto a polymeric iHILIC-(P) Classic HPLC column (HILICON, 200 x 2.1 mm; 5 µm), operated at a flow rate of 100 µl/min. A linear gradient (A: acetonitrile; B: 10 mM aqueous ammonium bicarbonate, supplemented with 0.1 µg/ml medronic acid) starting with 25% B and ramping up to 90% B in 14 minutes was used for separation. Using a TSQ Quantiva mass spectrometer (Thermo Fisher Scientific), the following SRM transitions were used for quantitation in the negative ion mode: *m/z* 87 to *m/z* 43 (pyruvate), *m/z* 89 to *m/z* 43 (lactate), *m/z* 117 to *m/z* 73 (succinate), *m/z* 124 to *m/z* 80 (taurine), *m/z* 133 to *m/z* 115 (malate), *m/z* 145 to *m/z* 101 (alpha ketoglutarate), *m/z* 167 to *m/z* 79 (phosphoenolpyruvate), *m/z* 169 to *m/z* 97 (dihydroxyacetone phosphate), *m/z* 173 to *m/z* 85 (aconitate), *m/z* 185 to *m/z* 79 (phosphoglycerate), *m/z* 191 to *m/z* 111 (citrate), *m/z* 259 to *m/z* 97 (hexose phosphates), *m/z* 339 to *m/z* 241 (fructose 1,6 bisphosphate), *m/z* 662 to *m/z* 540 (NAD), *m/z* 664 to *m/z* 407 (NADH), *m/z* 766 to *m/z* 408 (CoA), and *m/z* 808 to *m/z* 408 (Acetyl-CoA).

Amino acids were analyzed in a separate experiment using reversed phase chromatography. 100 µl of each extract was dried in a vacuum centrifuge and resolved in 100 µl of 0.1% formic acid in water. For each sample, 1 µl was injected onto a Kinetex (Phenomenex) C18 column (100 Å, 150 x 2.1 mm) connected with the respective guard column employing a 7-minute-long linear gradient from 99% A (1% acetonitrile, 0.1% formic acid in water) to 60% B (0.1% formic acid in acetonitrile) at a flow rate of 80 µl/min. Detection and quantification was done by LC-MS/MS, employing the SRM mode of a TSQ Altis mass spectrometer (Thermo Fisher Scientific), using the following transitions in the positive ion mode: *m/z* 90 to *m/z* 44 (alanine), *m/z* 106 to *m/z* 60 (serine), *m/z* 116 to *m/z* 70 (proline), *m/z* 118 to *m/z* 72 (valine), *m/z* 120 to *m/z* 74 (threonine), *m/z* 132 to *m/z* 86 (leucine and isoleucine), *m/z* 134 to *m/z* 74 (aspartic acid), *m/z* 148 to *m/z* 84 (glutamic acid), *m/z* 150 to *m/z* 133 (methionine), *m/z* 156 to *m/z* 110 (histidine), *m/z* 166 m/z to *m/z* 133 (phenylalanine), *m/z* 175 to *m/z* 70 (arginine), *m/z* 182 to *m/z* 136 (tyrosine) and *m/z* 205 to *m/z* 188 (tryptophan). Authentic standards were used for determining optimal collision energies of the SRM transitions and for validating experimental retention times via standard addition to a pooled quality control sample. The data interpretation was performed using TraceFinder (Thermo Fisher Scientific).

### Mitochondria isolation from zebrafish embryos

For mitochondrial isolation embryos were dechorionated using 1 mg/mL of pronase (Sigma-Aldrich). Dechorionated embryos were homogenized with a Dounce homogenizer (Sigma-Aldrich) in 1 ml mitochondria isolation buffer containing 210 mM mannitol, 70 mM sucrose, 1 mM EDTA, 10 mM HEPES (pH 7.5) supplemented with protease inhibitors cOmplete, EDTA-free protease inhibitor cocktail (Merck). The homogenate was cleared from nuclei at 500 g for 10 min 4 °C. Then, the supernatant was centrifuged at 9000 g for 10 min 4 °C to pellet mitochondria. Mitochondria were washed one time in 1 ml mitochondria isolation buffer. Protein concentration was determined with Bradford.

### Blue-native polyacrylamide gel and in gel activity staining

Blue-native PAGE (BNP) and in gel activity staining were performed as described before^82^ with minimal modifications. Isolated mitochondria were solubilized in BN-PAGE sample buffer (Thermo Fisher Scientific) with 8 g/g of digitonin on ice for 20 min. Every 100 μg of mitochondrial proteins were solubilized in 40 μl of buffer. The insoluble fraction was removed by centrifugation at 20.000 g for 20 min, and 40 μl of solubilized protein lysate was stained with 5 μl of 5% G-250 sample additive (Thermo Fisher Scientific). Wells of a Native-PAGE 3–12% Bis-Tris protein gel were washed with the dark blue cathode buffer, and 100 μg of mitochondrial proteins were loaded into each lane. The inner chamber was filled with the blue cathode buffer (Thermo Fisher Scientific), and the outer chamber with the native running buffer (Thermo Fisher Scientific). Protein complexes were separated by electrophoresis for 30 min at 150 V. After changing the buffer in the inner chamber to the light blue buffer, the gel was run for another 60 min at 250 V.

For measuring in-gel activity the protocol was slightly modified by using 200 ml of Native PAGE anode buffer (Thermo Fisher Scientific) supplemented with 0.022 g Coomassie Brilliant Blue G-250 instead of the regular light blue cathode buffer.

To measure the activity of respiratory complexes, gels were incubated in freshly prepared substrate solutions.

### Proteomics from isolated mitochondria

#### Proteomics and Phospho-proteomics from isolated mitochondria

*Sample preparation for label-free quantification of mitochondrial proteins* Mitochondrial pellets were lysed, and proteins were digested using the iST kit (Preomics) according to the manufacturer protocol. 3 µg of each peptide sample were analyzed by LC-MS/MS.

##### Sample preparation for TMT-labelled phosphopeptides

Isolated mitochondria were resuspended in 125 µl of 10 M urea, 50 mM HCl and incubated for 10 min at room temperature before the addition of 15 µl of 1 M Triethylammonium bicarbonate (TEAB) buffer pH 8. A Bradford assay was performed to determine the amount of protein. The lysates containing 300-500 µg of protein were supplemented with 20 mM DTT and 250 units Benzonase (250 U/µl, purity grade I, Merck) and incubated at 37°C for 1h. The alkylation was performed by adding 40 mM Iodoacetamide and incubating for 30 min in the dark. The reaction was quenched by addition of 10 mM DTT and incubation at room temperature for 30 min.

The samples were diluted to 6 M urea by addition of 100 mM TEAB and the proteins were digested with Lys-C (Wako) at an enzyme to protein ration of 1:50 for 3h at 37°C. Subsequently, the samples were further diluted to 2 M urea with 100 mM TEAB and trypsin (Trypsin Glod, Promega) was added at a ratio of 1:50 and the samples were incubated at 37°C over night. The samples were acidified by addition of Trifluoroacetic acid (TFA, Thermo Fisher Scientific, 28903) to reach pH 2. Peptides were desalted using C18 cartridges (Sep-Pak Vac 1cc (50mg), Waters). Peptides were eluted with 70% acetonitrile (ACN, VWR, 83639.320) and 0.1% formic acid (FA, Merck, 1.11670.1000), followed by freeze-drying. The peptide amount was determined by separating an aliquot of each sample on a capillary-flow LC-UV system using a PepSwift^TM^ Monolithic column (Thermo Fisher Scientific) and relating it to the peak area of 100 ng of Pierce HeLa protein digest standard (PN 88329; Thermo Fisher Scientific). 300 µg of tryptic peptides of 12 samples were dissolved in 100 µl 100 mM Hepes pH 7.6) and were labelled with a separate channel of TMTpro 16plex reagent (Thermo Fisher Scientific) according to the manufacturer’s description. Labelling efficiency was determined by LC-MS/MS with a small aliquot of each sample. Samples were mixed in equimolar amounts and equimolarity was again assessed by LC-MS/MS. The mixed sample was acidified to a pH below 2 with 10% TFA and desalted using C18 cartridges as described above.

Neutral pH fractionation was performed by applying a 30 min gradient from 4.5 to 45% ACN in 10 mM ammonium formate pH 7.5 on an UltiMate 3000 Dual LC nano-HPLC system (Dionex, Thermo Fisher Scientific) equipped with an XBridge Peptide BEH C18 (130 Å, 3.5 μm, 4.6 mm x 250 mm) column (Waters) (flow rate 1.0 ml/min). 40 fractions were collected and then pooled into 20 non-contiguous pools and quantified using a monolithic HPLC system.

A 10 µg aliquot was taken from each of these fractions for measurement of the proteome. Subsequently, the 20 fractions were further pooled into 10 pools, desalted via C18 cartridges as described above and used for phosphopeptide enrichment using TiO_2_.

An aliquot (peptide:TiO_2_ resin = 1:6) of TiO_2_ (Titansphere TiO_2_, GL Sciences, 5020-75000) was washed twice with 50% methanol (Fisher chemical, A456-212) and twice with glycolic acid solution (1 M glycolic acid (Sigma Aldrich, 124737-25G), 70% ACN, 3% TFA). Lyophilized peptides were dissolved in glycolic acid solution, mixed with the TiO_2_ resin and incubated for 30 min at room temperature with rotation. For washing and elution the bead suspension was transferred to Mobicol (MoBiTec) spin columns with a 10 µm pore filter at the outlet. The resin was washed twice with glycolic acid solution, twice with 200 μl 70% ACN and 3% TFA and twice with 1% ACN and 0.1% TFA. Phosphorylated peptides were eluted twice with 150 μl 300 mM ammonium hydroxide (VWR, 1.05432.1000). The eluates were combined, acidified to pH 2.5 with concentrated TFA, dried by vacuum concentration and dissolved in 20 µL of 0.1% TFA. 1 µg of each of the 20 nonenriched peptide pools and 100% of each of the 10 phosphoenriched pools were analyzed by LC-MSMS.

##### NanoLC-MS/MS analysis

The nano HPLC system (UltiMate 3000 RSLC nano system, Thermo Fisher Scientific) was coupled to a Q Exactive HF-X mass spectrometer equipped with a Proxeon nanospray source and, for the TMT experiment, to an Orbitrap Eclipse Tribrid mass spectrometer equipped with a FAIMS pro interface and a Nanospray Flex ion source (all parts Thermo Fisher Scientific).

Peptides were loaded onto a trap column (PepMap Acclaim C18, 5 mm × 300 μm ID, 5 μm particles, 100 Å pore size, Thermo Fisher Scientific) at a flow rate of 25 μl/min using 0.1% TFA as mobile phase. After loading, the trap column was switched in line with the analytical column (PepMap Acclaim C18, 500 mm × 75 μm ID, 2 μm, 100 Å, Thermo Fisher Scientific). Peptides were eluted at a flow rate of 230 nl/min, starting with mobile phases 98% A (0.1% formic acid in water) and 2% B (80% acetonitrile, 0.1% formic acid) and increasing linearly to 35% B over the next 180 min. This was followed by a steep gradient to 95% B in 5 min, held for 5 min and ramped down in 2 min to the initial conditions of 98% A and 2% B for equilibration at 30°C.

The Q Exactive HF-X mass spectrometer was operated in data-dependent mode using a full scan (m/z range 380-1500, nominal resolution 60,000, target value 1e6) followed by MS/MS scans of the 10 most abundant ions. MS/MS spectra were acquired using a normalized collision energy of 28, an isolation width of 1.0 m/z, a resolution of 30,000, a target value of 1e5 and a maximum fill time of 105 ms. Precursor ions selected for fragmentation (excluding charge states 1, 7, 8, >8) were placed on a dynamic exclusion list for 60 s. In addition, the minimum AGC target was set to 5e3 and the intensity threshold was calculated to be 4.8e4. The peptide match feature was set to preferred and the exclude isotopes feature was enabled.

The Eclipse was operated in data-dependent mode with a full scan (m/z range 350-1500, resolution 60,000, target 4e5) at 3 different compensation voltages (CV-40, -55,

-70) followed by MS/MS scans of the most abundant ions at a cycle time of 1.0 s per CV. MS/MS spectra were acquired using an isolation width of 0.7 m/z, target value of 1e5and intensity threshold of 2.5e4, maximum injection time of 120 ms, HCD with a collision energy of 34%, using the Orbitrap for detection, with a resolution of 50 k. For the detection of the TMT reporter ions, a fixed first mass of 110 m/z was set for the MS/MS scans. Precursor ions selected for fragmentation (including charge state 2-6) were excluded for 45 s. The Monoisotopic Precursor Selection (MIPS) mode was set to Peptide and the Exclude Isotopes feature was enabled.

##### Data Processing protocol

For peptide identification, the RAW files were loaded into Proteome Discoverer (version 2.5.0.400, Thermo Scientific). All MS/MS spectra were searched using MSAmanda v2.0.0.19924^83^.

For the protein identification set, the peptide mass tolerance was set to ±5 ppm and the fragment mass tolerance was set to ±15 ppm, with a maximum number of missed cleavages of 2, using tryptic enzyme specificity without proline restriction. Peptide and protein identification was carried out in two steps. For an initial search, RAW files were searched against the custom combined Uniprot/NCBI database named Danio_rerio.GRCz11.Uniprot_NCBI_21032019_nr99.fasta (58,524 sequences; 34,079,443 residues), supplemented with common contaminants and sequences of tagged proteins of interest using iodoacetamide derivative on cysteine as a fixed modification. The result was filtered to 1% FDR at the protein level using the Percolator algorithm^84^ integrated into Proteome Discoverer. A sub-database of proteins identified in this search was generated for further processing. For the second search, the RAW files were searched against the generated sub-database using the same settings as above and considering the following additional variable modifications: oxidation on methionine, deamidation on asparagine and glutamine, acetylation on lysine, phosphorylation on serine, threonine and tyrosine, methylation on lysine and arginine, di-methylation on lysine and arginine, tri-methylation on lysine, ubiquitinylation residue on lysine, biotinylation on lysine, formylation on lysine, glutamine to pyroglutamate conversion at peptide N-terminal glutamine and acetylation at protein N-terminus. Localization of post-translational modification sites within the peptides was performed using the ptmRS tool, based on the phosphoRS^85^. Identifications were again filtered to 1% FDR at the protein and PSM level, with an additional Amanda score cut-off of at least 150. Proteins were filtered to be identified by at least 2 PSMs in at least 1 sample. Peptides were subjected to label-free quantification using IMP-apQuant^86^. Proteins were quantified by summing unique and razor peptides and applying intensity-based absolute quantification^87^ (iBAQ). Protein abundance normalization was performed using sum normalization.

A direct search approach was used for the TMT analysis, with the following modifications to the workflow described above: The peptide and fragment mass tolerance were set to ±10 ppm. The ENSEMBL database used was Danio_rerio_GRCz11_ENSEMBL_104_pep_all.fasta (45,692 sequences; 26,804,937 residues). The phosphopeptide enriched and nonenriched fractions were separately searched using the following variable modifications : oxidation on methionine, deamidation on asparagine and glutamine, carbamylation on lysine, TMTpro 16plex tandem mass tag on lysine, carbamylation on peptide N-terminus, TMTpro 16plex tandem mass tag on peptide N-terminus, glutamine to pyro-glutamate conversion at peptide N-terminal glutamine, acetylation on protein N-terminus, phosphorylation on serine, threonine and tyrosine (only searched for the phosphopeptide enriched fractions). To allow a joint protein grouping, both search results were combined together in a common consensus workflow. Peptides were quantified based on reporter ion intensities extracted by the Reporter Ion Quantifier node implemented in Proteome Discoverer and proteins were quantified by summing unique and razor peptides.

To distinguish regulated phospho-peptides that are altered due to regulation at the phosphorylation site from changes due to regulation of the underlying protein, PSMs were separated into those acquired from phospho-enrichment and others acquired from the analysis of the non-enriched fractions to capture changes on the proteome. PSMs from the phospho enriched fractions were aggregated at the peptide isoform level. Other PSMs originating from non-enriched fractions were summarized at the protein level. These protein regulations were then used to correct the phospho-peptide regulations by the magnitude of the regulation of the corresponding protein. The corrected phospho-peptide regulations were used for further analysis.

### Microscopy

For live-imaging, dechorionated embryos were mounted in a drop of 0.8% low-melting agarose in PBS on round glass bottom dishes (Ibidi). Dishes were filled with E3 medium (5 mM NaCl, 0.17 mM KCl, 0.33 mM CaCl_2_, 0.33 mM MgSO_4_, 10^−5^% methylene blue). For imaging of mitochondria at early stages of embryogenesis, dechorionated embryos with EGFP-labeled mitochondria were fixed in 3.7% paraformaldehyde (PFA) in PBS overnight at 4 °C. Samples were washed in PBS and kept at 4 °C.

Images in Figure 5 were acquired on an inverted Olympus IX3 Series (IX83) equipped with a Yokogawa W1 spinning disk (SD) confocal scan head with a temperature of incubation of 29 °C for life imaging. An PlanApoX 100x/1.46 (Oil) WD 0.13 mm (Olympus) objective was used. Fluorophores were excited with corresponding lasers (488 nm for GFP, 561 nm for dsRED and 640 nm for Cy5, laser power was set to a minimum value allowing good signal-to-noise ratio and viability) and emitted signal was passed through a bandpass filter (525/50 for GFP and 617/73 for dsRED and 685/40 for Cy5) and detected using Hamamatsu Orca Flash 4.0 camera, at 700ms exposure and 480MHz readout.

The transplantation experiment multicolor data, has been corrected against chromatic shift using Huygens Professional (SVI). Correction was based on the multicolor bead data as a reference.

Images in Figure 6 were acquired as above, using double camera detection in order to avoid motion artefacts. The GFP signal was detected by camera 2 (Hamamatsu Orca Flash 4.0) with 525/50nm filter. The dsRED signal was detected by camera 1 (Hamamatsu Orca Flash 4.0) with 617/73nm filter.

### Imaging analysis

For assessing the mitochondria size and other parameters, different software packages were used. Preprocessing steps were done in Fiji, where movies were split into individual slices, a background subtraction was performed and images were converted to 8-bit, which was necessary for the next step, the segmentation via MitoSegNet^55^ (https://github.com/MitoSegNet/MitoS-segmentation-tool). Because the pretrained model provided with the software did not perform well enough, the model was re-trained and adapted to our own dataset. QuPath^88^ and its annotation tools were used to create ground-truth images which were then used to update the deep learning model. Using this model, binary masks were created of our 2D slices, which were analyzed in Fiji.

### Transplantation experiment

Donor and host embryos were manually dechorionated with watchmaker forceps in 0.3 x Danieau’s medium (17.4 mM NaCl, 210 µM KCl, 1.5 mM Hepes buffer (pH 7.1-7.3), 180 µM Ca(NO_3_)_2_ and 120 µM MgSO4). A small amount of cytoplasm from Tg(*actin::mts-EGFP*) embryos was transferred into Tg(*actin::mts-dsRed*) embryos using a bevelled borosilicate needle (20 μm inner diameter with spike; Biomedical Instruments) connected to a syringe system mounted on a micromanipulator. Embryos were incubated at 28 °C for 1 h to recover before imaging on the inverted Olympus IX3 Series (IX83).

### Electron microscopy

Zebrafish oocytes were collected at different timepoints post fertilization and immediately fixed in 2.5% glutaraldehyde (EM-grade; Agar Scientific, UK) in 0.1 M cacodylate buffer over night at RT. After washing three times in 0.1 M cacodylate buffer, oocytes were post-fixed in 1% osmium tetroxide (EMS, USA) in 0.1 M cacodylate buffer on ice for another hour. Samples were rinsed in 0.1 M cacodylate buffer for three times 10 min each. Oocytes were dehydrated in a graded series of acetones: 40%, 60%, 80%, two times 95% and three times 100%; each step was performed for 15min on ice. Subsequently the samples were infiltrated with Agar 100 epoxy resin (Agar Scientific, UK) and polymerized at 60 °C for 48 h.

Resin blocks were cut using a Leica EM UCT ultramicrotome (Leica Microsystem, Austria) with a nominal section thickness of 70 nm and picked up onto hexagonal Cu/Pd grids with a mesh size of 100 lines per inch, previously coated with a self-made Formvar® film (both grids and Formvar powder from Agar Scientific, UK). To enhance contrast, grids were post-stained with 2% aqueous uranyl acetate at pH 4 (Merck, Germany) and Reynold’s lead citrate. Grids were inspected in a FEI Morgagni 268D transmission electron microscope (Thermo Fisher Scientific, The Netherlands), operated at 80 kV. Examined regions were chosen randomly. Digital images were taken on a Mega View III CCD camera from Olympus-SIS (Germany).

### Proximity ligation assay (PLA)

PLA was performed according to the Duolink In Situ Red Starter Kit Mouse/Rabbit (Sigma-Aldrich, DUO92101) protocol with some modifications. Dechorionated samples were fixed with 3.7% FA in PBS at 4 °C overnight. Fixed samples were washed with PBS-T buffer (0.1% Tween20 in 1X PBS) five times for 5 min at room temperature. Next, samples were permeabilized with PBS-Tr (0.5% Triton X-100 in PBS) for 30 min at RT and then blocked with blocking solution (1% DMSO, 2% BSA, 5% NGS in PBS-Tr (0.1%)) overnight at 4 °C. All the following steps, including primary antibody incubation, PLA probe incubation, ligation, amplification, and final washes, were performed according to the protocol provided with the kit except that primary antibodies were diluted 1:100 in blocking solution (1% DMSO, 2% BSA, 5% NGS in PBS-Tr (0.1%)). Primary antibody pairs used include mouse anti-Hspa9 (Santa Cruz Biotechnology, sc-133137), rabbit anti-SigmaR1 (Proteintech, 15168-1-AP) and rabbit anti-Flag (Cell Signaling, 14793).

### RNA-seq

Embryos were collected as soon as they were laid and put into E3 medium (5 mM NaCl, 0.17 mM KCl, 0.33 mM CaCl_2_, 0.33 mM MgSO_4_, 10^−5^% methylene blue) containing inhibitors. Embryos were kept in this medium for the duration of the experiment. For this experiment we used already established concentrations for inhibitors: oligomycin 1 µM (Millipore, 495455), actinomycin D 10 µg/ml (Sigma-Aldrich, A1410). Untreated embryos were kept alongside as controls. Total RNA was extracted using the RNeasy Mini Kit (Qiagen, 175023525). Samples were submitted to the VBCF NGS facility for Quantseq library preparation using the Quantseq kit (Lexogen), followed by sequencing using the NextSeq2000 P2 SR100 platform (Illumina). Reads were pre-processed using umi2index (Lexogen) to add the UMI sequence to the read identifier, and trimmed using BBDuk v38.06 (ref=polyA.fa.gz,truseq.fa.gz k=13 ktrim=r useshortkmers=t mink=5 qtrim=r trimq=10 minlength=20). Reads mapping to abundant sequences (mitochondrial chromosome, phiX174 genome, and silva zebrafish rRNAs from the SILVA rRNA database) were removed using bowtie2 v2.3.4.1 alignment. Remaining reads were analyzed using genome and gene annotation for the GRCz11 assembly obtained from *Danio rerio* Ensembl release 104. Reads were aligned to the genome using star v2.6.0c, alignments were processed using collapse_UMI_bam (Lexogen) and reads in genes were counted with featureCounts (subread v1.6.2) using strand-specific read counting (-s 1).

### Experimental design and statistical analysis

All experiments reported in this study were performed at least twice. Zebrafish were categorized by genotype and age, with adult males and females used to generate embryos ranging in age from 3 months to 1 year. *In vivo* samples were randomly assigned to experiments and treated equally throughout. Sample size was not statistically predetermined. Fish and embryos were randomly selected for *in vivo* experiments.

For Figure 4, proteins with a minimum of 3 unique peptides in 2 samples of 1 condition, or a minimum of 2 unique peptides in 2 samples of 2 conditions, were retained for further analysis. For those, significant changes in protein abundance between timepoints were evaluated with limma v3.58.1 using functions lmFit for linear model fitting and eBayes for moderated t-statistics computation. Significant proteins were selected at an adjusted P-value threshold of 0.05 and log2FC threshold of 1.

Statistical analyses and graphs were performed using R statistical software (version 2023.12.1). Standard statistical tests were selected to account for sample normality, and post hoc tests (Benjamini & Hochberg correction) were used for multiple comparisons. Details of the tests performed, and corresponding P-values are provided in the graphs and figure legends.

In the Figures 1, 5, 6 and S5, the geom_smooth() function from the ggplot2 package in R was used to fit a locally estimated scatterplot smoothing (LOESS) curve. This nonparametric regression method was used to highlight the trend in the data without assuming a specific model.

### Data availability

RNA-seq data generated in this study is available at GEO with the accession numbers GSE267318 (Quant-seq). Mass spectrometry proteomics data has been deposited to the ProteomeXchange Consortium via the PRIDE (30395289) partner repository with the dataset identifier PXD049073 (mitochondrial proteomics) and PXD051555 (mitochondrial phospho-proteomics).

